# Maternal exercise during lactation reprograms obesity-related changes in mammary metabolism to optimize milk fatty acids and offspring energy expenditure

**DOI:** 10.1101/2025.09.04.674229

**Authors:** Gertrude Kyere-Davies, Kaitlyn B. Hill, Gregory P. Mullen, Rohan R. Varshney, Snehasis Das, Alexandrea Martinez, Jacob W. Farriester, Michael Kinter, Cassie M. Mitchell, Kevin R. Short, Elvira M. Isganaitis, David A. Fields, Michael C. Rudolph

## Abstract

Maternal obesity alters breast milk composition in ways that may predispose infants to excess adiposity. While maternal exercise during lactation has been associated with favorable shifts in milk metabolites in humans, the mechanisms by which exercise remodels the mammary gland and milk lipid profile to influence offspring metabolism remain unclear. We developed a mouse model incorporating daily moderate treadmill exercise during lactation, indirect calorimetry, stable isotope tracer respirometry, and mammary epithelial cell (MEC) proteomics in lean (LN) and diet-induced obese (OB) dams. Maternal obesity broadly remodeled the MEC proteome, reducing enzymes of *de novo* fatty acid synthesis and altering lipid transport and oxidative pathways. These molecular adaptations corresponded to higher milk triglyceride content and shifts in fatty acid composition, including an elevated omega-6 to omega-3 fatty acid ratio. The exercise (EX) intervention during lactation reset MEC protein networks, enhancing translational and vesicle transport pathways while reducing fatty acid desaturation, relative to the sedentary (SED) group. In OB dams, exercise increased milk medium-chain fatty acid (MCFA) levels and partially corrected the n6/n3 FA ratio. Offspring nursed by OB-EX dams exhibited higher whole-body energy expenditure, increased fatty acid oxidation, and improved metabolic flexibility compared to litters consuming OB-SED milk. Together, maternal exercise during lactation remodels mammary metabolism and milk fatty acid composition in obese dams, enhancing neonatal lipid oxidation and energy expenditure. These findings highlight lactation as a modifiable window, wherein maternal activity influences milk composition and infant metabolic health.

**New and noteworthy:** Maternal obesity alters milk fatty acid composition, with consequences for infant metabolism. Exercise during lactation in obese dams remodeled the mammary epithelial cell proteome, increasing medium-chain fatty acids in milk and enhancing lipid oxidation and energy expenditure in offspring.

## 1.0 INTRODUCTION

Lactation is a critical biological event that supports maternal and infant health, offering both immediate and long-term benefits to mother and child. The American Academy of Pediatrics and the World Health Organization recommend exclusive breastfeeding for the first six months of life, with continued breastfeeding up to two years alongside complementary foods (1). Breast milk provides all essential macro- and micronutrients, along with numerous bioactive compounds that sustain infant growth, development, and immune function (1–3). Milk composition is dynamic, with the compositional make-up changing over the course of lactation, in response to maternal physiology, and even within a single feeding session (2, 4). For infants, breastfeeding lowers their risks for contracting infectious diseases (5–8), experiencing sudden infant death syndrome (9), and developing chronic conditions such as type 2 diabetes, asthma, and certain childhood cancers (1, 10–12). Mothers benefit as well, since prolonged lactation is associated with lower incidence of breast, cervical, and ovarian cancers, and reduced risk of metabolic diseases including type 2 diabetes (13–15). These examples emphasize lactation as a unique window of opportunity to improve the long-term health trajectory of both mother and infant. Given its dual role in supporting infant development and maternal metabolic recovery, increasing attention in the field has been directed toward how maternal physiological states, such as nutritional status, adiposity, and physical activity, coordinate milk macronutrient, micronutrient, and bioactive compositions and, in turn, how variance in milk composition shapes offspring health outcomes.

Although there are several established benefits of breastfeeding, not all mothers are able to initiate or sustain lactation, particularly those with underlying metabolic conditions such as obesity. Lactation insufficiency, defined as an inability to successfully initiate and/or maintain lactation, affects about 20% of women globally and up to 50% of women in the U.S., making it a predominant cause for early breastfeeding cessation (16). Maternal obesity has been associated with an increased risk for lactation challenges, potentially limiting infants’ access to the full metabolic and immunologic benefits of breastfeeding (17, 18). These difficulties are believed to arise, in part, from disruptions in the hormonal regulation of mammary gland development, milk synthesis, and milk secretion (19, 20). Animal models provide mechanistic insight into this relationship, wherein maternal obesity in rodents increases the risk of primary lactation failure and neonatal mortality, while overfeeding in dairy cows can induce “fat cow syndrome,” a metabolic condition linked to disrupted lactogenesis (20). In human studies, maternal pre-pregnancy BMI ≥ 30.0 kg/m^2^ is associated with delayed initiation of lactation (21). Collectively, these observations suggest that maternal OB may alter mammary gland function through complex mammary developmental and/or physiological mechanisms that are still being actively investigated.

In addition to influencing lactation success, maternal OB is associated with alterations in breast milk composition that may contribute to an obesogenic postnatal environment, potentially increasing the infant’s risk for excess adiposity and metabolic disease later in life (22). For example, we and others have shown in rodent models that the fatty acid composition of mother’s milk, specifically high omega-6 to omega-3 (n6/n3) fatty acid ratio, affects the fat mass accumulation and favor nutrient storage over oxidation as well promote pro-inflammatory and oxidative stress in the offspring (23–25). These findings highlight the importance of balanced milk n6/n3 FA composition in shaping early adipocyte metabolic programming. The clinical relevance of these findings is tied to the observation that people in the U.S have an elevated n6/n3 ratio in their diets (26, 27). Conversely, maternal OB is associated with a decrease in medium chain fatty acids (MCFAs) such as lauric acid and increase in sugars and amino acids in the milk, which can affect infant metabolic programming (28). MCFAs are present in breastmilk and present as important metabolic fuels for infant growth and development (29).

Compounding the effects of FA exposures, metabolic phenotype during the perinatal period, including obesity, gestational diabetes, and type-2-diabetes, has been linked to increased risk of childhood obesity in both human and animal studies (30). For example, higher maternal BMI during pregnancy and lactation is associated with greater infant fat mass accumulation and increased susceptibility to early weight gain (31), underscoring the need for targeted interventions during this critical window. Notably, recent findings suggest that maternal physical activity can influence the milk metabolome, with specific metabolites and lipids associated with infant adiposity trajectories (32, 33). Together, these findings emphasize the potential to harness maternal lifestyle factors during lactation to support infant energy metabolism and mitigate long-term metabolic risk.

Most interventions targeting lactation have emphasized education and psychological support, yet few have explored how maternal physiological adaptations can be leveraged to improve lactation outcomes. In this study, we examine the potential for maternal exercise, a modifiable lifestyle factor, to mitigate the lactational consequences due to maternal diet induced obesity that influence both milk quality and offspring metabolic programming. Exercise is well-established in the prevention and management of metabolic diseases, including obesity, type 2 diabetes, and cardiovascular conditions (34). In humans, physical activity during pregnancy is associated with reduced risk of gestational hypertension, preeclampsia, and gestational diabetes (35–37). Animal models further demonstrate that maternal exercise during gestation can enhance insulin sensitivity and glucose homeostasis in offspring across the lifespan (38, 39). More recently, exercise during lactation has gained attention for its ability to modulate milk composition. We found that an acute bout of maternal exercise at 1-month postpartum increased breast milk levels of the lipokine 12,13-diHOME in humans (33). This metabolite has been associated with enhanced fatty acid uptake and oxidation in adult tissues (40), and in our cohort, was associated with lower infant adiposity and improved weight-for-length z-scores suggesting that maternal exercise can enrich breast milk with anti-obesogenic lipids (33).

However, whether maternal exercise during lactation alone, independent of gestational exposures, is sufficient to reprogram postnatal metabolism remains unknown. Most animal studies combine maternal exercise during gestation and lactation, whereas our study carefully isolates exercise to the lactation period. This design allows us to assess the main effects of both maternal obesity and daily moderate treadmill exercise in lactating dams. Additionally, by providing a cohort of dams with a diet rich in linoleic acid, we examine how maternal dietary fatty acid composition interacts with exercise to influence maternal energetics, milk composition, and offspring metabolic phenotype. Our findings offer mechanistic insights into how lactation-stage interventions can modify mammary gene expression and enzymatic pathways of milk production, enhancing the milk fatty acid composition consumed by pups. These adaptations, in turn, have the potential to correct offspring metabolic fuel preferences and potentially mitigate childhood obesity risk.

## 2.0 MATERIALS AND METHODS

### 2.1 Animal care and feeding protocol

Seven (7) week C57BL/6J female mice were obtained from The Jackson Laboratory (Bar Harbor, Maine) and maintained in the Research Barrier Facility, the Division for Comparative Medicine, the University of Oklahoma-Health Campus (OUHC). OUHC Institutional animal Care and Use Committee (IACUC) approved all animal procedures and housing conditions. Mice acclimatized for 7-days in our mouse metabolic phenotyping satellite room which is maintained at 26°C, until they reached 8-weeks of age. After acclimatization, the animals were randomized to control diet (D20071402, 15% kcal soybean fat, 65% kcal corn starch and dextrose; Research Diets, New Brunswick, NJ, USA) or an obesogenic, high-fat high-sucrose (HFHS) diet (D21101202, 45% kcal corn oil fat, 35% kcal primarily sucrose, with some corn starch and dextrose) for 10-weeks to induce an obese maternal phenotype (>26% body fat by quantitative magnetic resonance, (qMR; Echo MRI Whole Body Composition Analyzer; Echo Medical Systems, Houston, TX, USA) (23, 41), after which all dams were paired for mating. Mice provided the control diet were designated “LN dams,” and those on the HFHS diet were in the “OB dams” group. At 18 weeks old, the dams in both LN and OB maternal groups were mated with age-matched C57/B6J male mice, allowed to undergo normal gestation, parturition, and lactation (*Fig. 1A*) (24)

**Figure 1:**
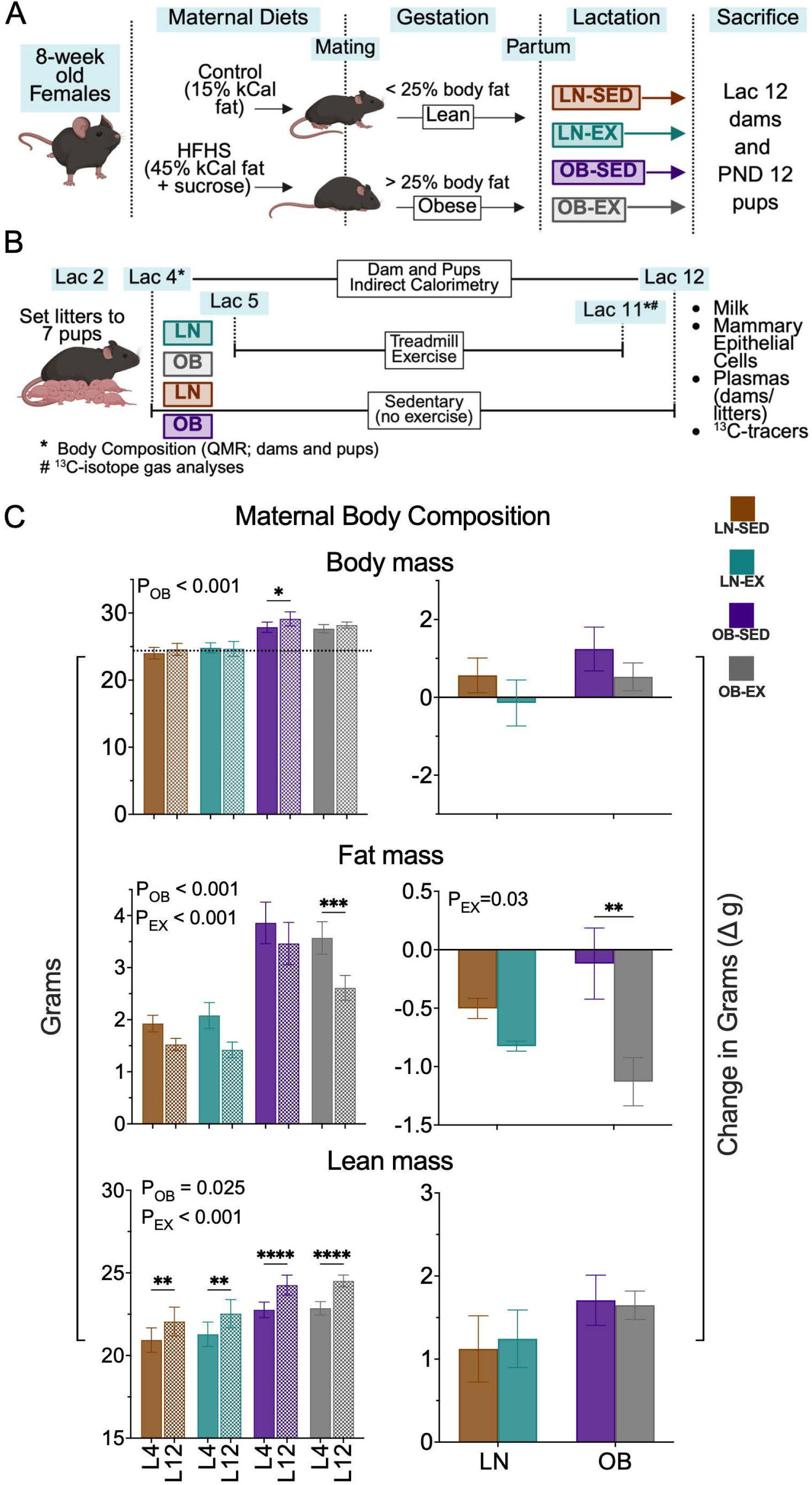
Maternal exercise increases loss of fat mass in obese lactating dams. The effects of maternal exercise on body composition, expressed in grams, of lactating dams were evaluated before and after a 7-day exercise period. (A) Experimental model for feeding and generating groups. (B) Experimental design for events and timelines. (C) Body mass, fat mass, and lean mass of dams (n=5 LN-SED/EX, n=10 OB-SED and n=13 OB-EX) were measured using EchoMRI on lactation day 4 (Lac4) before the exercise beginning on Lac5. Post-exercise data were collected on Lac12 after the exercise concluded on Lac11. Delta data were calculated by subtracting pre-exercise measurements from post-exercise measurements. Data are presented as means ± SEM, and a 2-way ANOVA determines statistical significance for the main effects of obesity and exercise on the data denoted *P_OB_* or *P_EX_*. Asterisks represent comparison between Lac4 and Lac12, and within the same group and denoted by *\*p<0.05, **p<0.01, ***p<0.001, ****p<0.0001*. Lac = Lactation day; PND = post-natal day; LN = lean; OB = obese; SED = sedentary; EX = exercise.

### 2.2 Experimental design

#### Body composition and Indirect Calorimetry analysis

On lactation day 2 (Lac2, 2 days after parturition), pups were standardized to 7-8 pups/dam, and dams from LN and OB groups were randomized into exercise (EX) and sedentary (SED) groups, giving four experimental conditions (LN-EX, LN-SED, OB-EX and OB-SED). On Lac4, pre-exercise body composition was measured separately for the dams and their respective litters (dams were measured alone, whiles litters were measured as one unit for each dam) using qMR. The dam-litter dyads were then placed into indirect calorimetry chambers (IDC, Sable Systems International, Las Vegas, Nevada) for metabolic phenotyping from Lac4-12 (41) (*Fig. 1B*). Dam metabolic phenotype measures included energy intake (EI), energy expenditure (EE), energy balance (EB), locomotion, and respiratory exchange ratio (RER). From the consumed O2 and expired CO2, carbohydrate and lipid oxidation were estimated using the equations 1.695xVO2 – 1.701xVCO2 and 4.585xVO2 – 3.226xVCO2, respectively. Dam measurements were taken with litters present in the cages. Litter metabolic phenotype included RER, EE, and systemic nutrient oxidation via stable isotope gas analysis. Litter measurements were taken with dams separated from the pups, by removing dams from the cages for about 3 to 4 hours on Lac11 only.

#### Exercise regimen and Stable Isotope Gas Analysis

From Lac5 - Lac11, the EX cohort underwent a controlled (involuntary) treadmill exercise using a 5 min warm-up at 6 m/min, moderate exercise at 12 m/min for 40 min, and a cool-down at 6 m/min for 5 min schedule. Dams were removed from the cages to run on the treadmill every morning at 9:00 am. Stable isotope tracers were used to quantify substrates of fuel utilization for both dams and pups on Lac11 following the final bout of maternal exercise, according to our methods (24). Briefly, dams were separated from their litters and administered a uniformly labeled stable ^13^C_6_ glucose (dissolved in saline) by intraperitoneal (IP) injection, and systemic glucose oxidation was quantified by measuring the exhaled ^13^CO_2_ per minute using a stable isotope gas analyzer (Sable Systems International). Separately, individual litters of dams from each group were administered 100mg/kg of uniformly labeled ^13^C_16_-palmitate orally (∼500ug/pup dissolved in 50μL total vol. of peanut oil) to quantify systemic fatty acid oxidation. On Lac12, post-exercise body composition was measured separately for both dams and litters prior to sacrifice, and tissues and samples were collected for the downstream assays. Blood samples from dams were collected via cardiac puncture while samples from litters were collected via decapitation and exsanguination and pooled together. Plasma was separated from whole blood by centrifugation at 1,000 × g for 10 min at room temperature and stored at -80°C until needed.

### 2.3 Mouse milk collection

Milk was collected on Lac12 as previously reported (42). Dams were separated from their litters for 2 - 4 hours before the milking was conducted to ensure adequate milk accumulation. Following separation, dams were immobilized using IP injection of Xylazine 5-8 mg/Kg (in sterile saline, 100 µL). Once the Xylazine had taken effect, 0.2 units of Oxytocin in sterile saline was administered IP to the dams to induce milk let down. Milk was expressed from all glands by using a hand-held vacuum apparatus connected to 1.5 mL microfuge tube, briefly collected by table-top centrifuge ‘sweat-spin,’ and stored immediately at -80°C for later assays (43).

### 2.4 Milk composition and nutrient density analysis

Total milk lipid, protein, and lactose levels were quantified by colorimetric assays according to the manufacturer protocols. Frozen milk samples were incubated at 37°C to thaw and reduce viscosity for the assays. Five (5) µL of milk was diluted into 125µL of potassium phosphate buffer pH 6.8. The suspension was centrifuged at 13,000 x g at room temperature for 3 min to clear the milk. Total protein was quantified using the Pierce™ BCA Protein Assay Kit (Thermo Scientific 23227). Lactose was quantified using the Sigma Lactose Assay kit (Sigma-Aldrich #MAK487). Total milk triglyceride was quantified using the Triglyceride Reagent Set kit (HORIBA Instruments #T7532-120).

### 2.5 Mammary epithelial cells isolation from whole mammary gland

Mammary epithelial cells (MECs) were isolated using the previously described protocol (44), after lactating dams had been euthanized using carbon dioxide and cervical dislocation according to IACUC approval. MECs were isolated from dams that had not previously been milked.

### 2.6 RNA isolation and quantitative real-time PCR (qPCR)

#### RNA isolation

MECs stored in RLT buffer were homogenized for RNA extraction using Bead Mill 24 Homogenizer (Fisherbrand™, 15-340-163). Total RNA was extracted using the RNeasy® Plus Mini Kit (Qiagen GmbH, Hilden, Germany, 74134) according to the manufacturer’s protocol. The Nanodrop 8000 Spectrophotometer (Thermo Scientific, ND-8000-GL) was used to quantify the concentration of total RNA and the integrity was assessed using the 4200 TapeStation Systems (Agilent Technologies, G2991BA). For quantitative real time PCR analysis, 500ng of total RNA was reverse transcribed into cDNA using the Verso cDNA Synthesis Kit as described in the manufacturer’s protocol, using only oligo-dT as the primer (ThermoFisher, MA, USA). cDNA representing 25ng of total RNA was added to each qPCR reaction containing TaqMan Fast Advanced Master Mix and TaqMan primers specific to Acly, Acc1, Fasn, Srebp1, Olah, Socs1 and 3, Spot14 (Thrsp), Csn2, Wap, Lalba, Btn and Xdh for amplification (ThermoFisher, MA, USA). RPL32 was the reference gene.

### 2.7 Protein extraction and proteomics analysis

#### Protein extraction

MECs in MG lysis buffer were homogenized using the Bead Mill 24 Homogenizer (Fisherbrand™, 15-340-163) for protein extraction. Total protein concentration was estimated using the Pierce™ BCA Protein Assay Kit (Thermo Scientific 23227).

#### Sample preparation for proteomics

Protein samples were transferred into a microfuge tube. 150 µL 1% SDS and 20 µL BSA internal standard were then mixed with the samples, heated and precipitated overnight using acetone. A final concentration of 1 µg/µL Laemmli buffer was added to the precipitated samples and 20 µL (a total of 20µg protein) of each of the samples were run in a 1.5cm SDS-PAGE gel. The lanes were cut as individual samples, washed, reduced, alkylated, and digested with 1 µg of trypsin overnight. The peptides were extracted from the gel with 50% acetonitrile, taken to dryness by Speedvac, and reconstituted in 200µL 1% acetic acid for analysis.

#### Data independent acquisition (DIA)

The Q Extractive plus instrument (ThermoScientific) was used in DIA mode with a 20m/z window working from m/z 350 to 950. The orbitrap was operated at a resolution of 17,500. A full scan spectrum at a resolution of 70,000 was acquired each cycle giving 7-8 data points across a 30 sec chromatographic peaks. The data were analyzed using the program DIANN, which uses neural network methods to analyze the DIA datasets versus the respective Uniprot proteome database from Embl. Linear Models Microarray and RNA-seq Data (LIMMA) was used to generate differentially enriched proteins and pathway analysis was performed using the Enrichr web application.

### 2.8 Lipid gas chromatography-mass spectrometry (GC/MS)

Lipids from milk and plasma samples were extracted and fatty acid profiles analyzed by GC/MS (23). Five (5) µL of milk or 6 µL of plasma was added to 500 µL potassium phosphate buffer pH 6.8. 1N prewashed HCL (40 µL for milk and 10 µL for plasma) was added and swirled lightly to acidify the samples. Then 500 µL of methanol was added to the solution and vortexed briefly. Total lipids were extracted by adding 1 mL 3:1 (v/v) isooctane/ethyl acetate solution twice and 1 mL hexane once. The organic layers collected from each extraction were pooled and dried using nitrogen. Dried lipids were resuspended in Isooctane (300 µL for plasma and 500 µL for milk). To quantify the non-esterified fatty acid (NEFA) fraction, 100 µL of the resuspended lipids from the plasma or 20 µL from the milk samples were transferred into a fresh tube and 25ng of blended FA stable isotope internal standard was added (23), brought to dryness, derivatized by adding 20 µL Pentafluorobenzyl bromide (PFBBr) and 40 µL N, N-Diisopropylethylamine (DIEA) and incubated for 25 minutes at 37°C. The derivatized samples were then dried and resuspended in 100 µL of hexane for injection into the GC/MS for analysis. For the total fatty acid (TFA) fraction, 50 µL of the resuspended lipids (plasma) or 20 µL (milk) samples were transferred into a fresh tube and 66.7 ng (plasma) or 500 ng (milk) of blended FA stable isotope internal standard was added. The samples were dried, then resuspended in 500 µL of 100% ethanol and saponified with 1N NaOH (500 µL each) at 90°C for 30 min, followed by neutralization with 525 µL 1N HCl. The saponified fatty acids were then extracted twice with 1.5 mL hexane, dried, derivatized as above, dried again. Plasma samples were resuspended in 267 µL hexane for injection into the GC/MS. Milk samples were resuspended in 500 µL hexane and diluted 4x before injection into the GC/MS. For both the NEFA and TFA fractions, 1 µL of pentafluorobenzyl-fatty acid derivatives was injected and data were collected by GC-MS running in NCI mode (8890 GC, 5977B MSD, Agilent) using the DB-1ms UI column (122-0112UI, Agilent) with the following run program: injector temp was 350°C, 80°C hold for 3 minute, 30°C/minute ramp to 125°C, no hold, 25°C/minute ramp to 320°C and hold for 2 minutes. The flow for the methane carrier gas was set at 1.5mL/minute. Fatty acids with chain lengths from 8 to 22 carbons were identified using full scan negative chemical ionization mode. Peak areas of both standard and samples were measured, and the ratio of each analyte’s ion signal to that of the internal standard was calculated to acquire the data (24).

### 2.9 Statistical analysis

Results are presented as mean ± SEM, with *p* values less than 0.05 considered significant. Two-way ANOVA was performed for analyses of differences between groups as noted in the figure legends. Tukey post-hoc multiple testing comparisons were performed where appropriate using GraphPad Prism Software, Version 10.6.0.

## 3.0 RESULTS

### 3.1 Maternal exercise during lactation decreases fat mass without altering total weight

We determined the exercise effect on body weight and body composition during the lactation window by measuring LN and OB dams before and after the exercise intervention (*Fig. 1B*). Maternal exercise had no significant effect on total weight in either the LN or OB dams. As designed, the main effect on maternal body weight was driven by obesity (*Fig. 1C*, *P*_OB_<0.001). OB dams weighed more than the LN dams before (Lac4, *P*_OB_=0.0003, Supplemental Figure S1A) and after the exercise intervention (Lac12, *P*_OB_=0.0001, Supplemental Figure S1B). No significant loss of body weight was observed with our moderate regimen of maternal exercise, however, OB-SED dams gained weight by Lac12 (*p*=0.01). While the OB dams had nearly twice the fat mass of the LN dams at both Lac4 and Lac12 (*Fig. 1C*, *P*_OB_<0.001), maternal fat mass was significantly reduced as a main effect of EX (*Fig. 1C*, *P*_EX_<0.001). The change in fat mass from Lac4 to Lac12 was significantly affected by maternal exercise (*P*_EX_=0.03), with the largest decrease observed in the OB-EX dams (*p*=0.003).OB dams increased their lean mass at Lac12 relative to Lac4, (*Fig. 1C*, *P*_OB_=0.025), while maternal exercise significantly increased overall gain of lean mass (P_EX_<0.001). However, no difference was observed in the change in lean mass between Lac4 to Lac12 for all groups (*Fig. 1C*). These findings indicate that maternal exercise during lactation selectively limited fat mass accrual in obese dams, without compromising gain of lean mass total body weight maintenance.

OB dams had significantly greater EI (*Fig. 2A*, *P*_OB_=0.009), especially for the OB-EX group relative to the OB-SED dams (*p*=0.05). Cumulative and total EE (*Fig. 2B*) was unchanged by maternal exercise, not accounting for the energetic cost of the exercise bout. Interestingly, the LN-EX dams tended to reduce their EE (*Fig. 2B*, *p*=0.09) relative to LN-SED, while EE in the OB group was unchanged. The OB dams reached greater positive energy balance than the LN dams (*P*_OB_=0.005), particularly in the OB-EX group relative to the OB-SED group (*p*=0.04), likely resulting from their increased in energy intake (*Fig. 2C*). Together, metabolic phenotyping data indicate that OB-EX dams responded to maternal exercise by increasing food intake without changing EE, while conversely, the LN-EX dams decreased EE without changing their food intake. These differences in body composition and systemic metabolic responses suggest different maternal adaptation strategies for lean and obese dams to the metabolic demand of lactation during moderate exercise.

**Figure 2:**
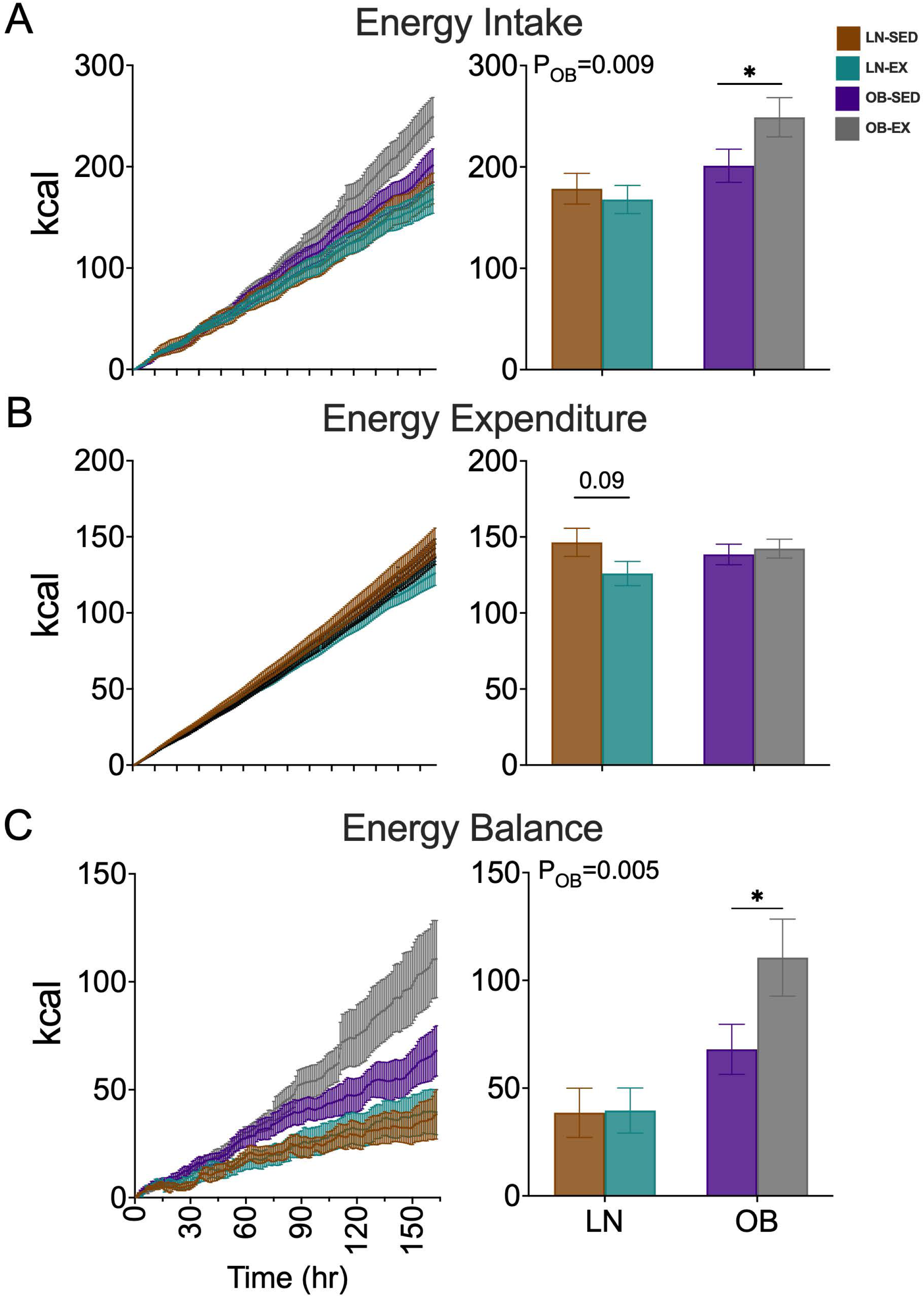
Maternal exercise differentially affects eating habits of lactating dams based on maternal phenotype. Maternal metabolic phenotype was assessed using metabolic indirect calorimetry (IDC). The effects of maternal exercise on whole-animal metabolism were determined by measuring (A) cumulative and total energy intake, (B) cumulative and total energy expenditure, and (C) cumulative and total energy balance (n=5 LN EX/SED, n=6 OB-SED, n=8 OB-EX). Dams were placed in individual metabolic cages with their litters from Lac4 for acclimatization before exercise. Primary data was collected on Lac5 to Lac11. Data are presented as means ± SEM, and a 2-way ANOVA determines statistical significance for the main effects of obesity and exercise on the total data denoted P_OB_ or P_EX_. Asterisks represent comparison within the same group and denoted by *\*p<0.05, **p<0.01, ***p<0.001, ****p<0.0001*.

### 3.2 Lean and obese dams adopt distinct systemic metabolic strategies in response to exercise

The respiratory exchange ratio (RER; VCO_2_/VO_2_) indicates the general metabolic fuel utilization needed by lactating dams. Dams in the OB group had a significantly lower average RER relative to LN dams (*Fig. 3A*, *P_OB_*<0.0001), indicating a shift away from carbohydrates as a fuel for nutrient oxidation, likely due to the 45% kCal fat present in the diet of this group. The main effect of EX significantly increased RER (*Fig. 3A*, *P_EX_*=0.03), indicating that maternal exercise shifted fuel utilization similarly despite vastly different dietary compositions. A main effect of OB was observed, with OB dams having lower carbohydrate oxidation than LN dams (*Fig. 3B*, *P_OB_*=0.003), even though the OB maternal diet was laden with sucrose. Conversely, a main effect of EX for lipid oxidation was seen (*Fig. 3B*, *P_EX_*=0.02), particularly in LN-EX dams, although greater overall lipid oxidation was prevalent in the OB group (*P_OB_*=0.0002). The lower carbohydrate oxidation in the OB dams suggested a possible reduction in systemic glucose metabolism, which is needed, in part, for *de novo* fatty acid synthesis. To ascertain how dams metabolized glucose, an independent cohort was administered uniformly labeled ^13^C_6_-glucose (2g/kg body weight) by IP injection, after which the ^13^CO_2_ emitted in the dams’ breath was quantified by stable gas isotope analysis (24). OB dams respired significantly more ^13^CO_2_ than LN dams with area under the curve (AUC) analysis (*Fig. 3C*, *P_OB_*<0.0001), and there was a main effect of EX on increasing ^13^C-glucose oxidation (*P_EX_*<0.0001). Interestingly, there was an interaction in OB-EX dams, which had significantly greater ^13^CO_2_ respiration (*P_INT_*<0.0001). There was an effect of maternal EX on the highest amount of glucose oxidized where the SED dams tended to have a higher peak of oxidation than EX-dams (*Fig. 3C*, *P_EX_*=0.08, inset). However, EX-dams had slower but more sustained oxidation, resulting in an overall increase, irrespective of weight status. This suggests a metabolic priority in biosynthetic pathways, such as those that support milk fatty acids, over utilization of substrates for energy production.

**Figure 3:**
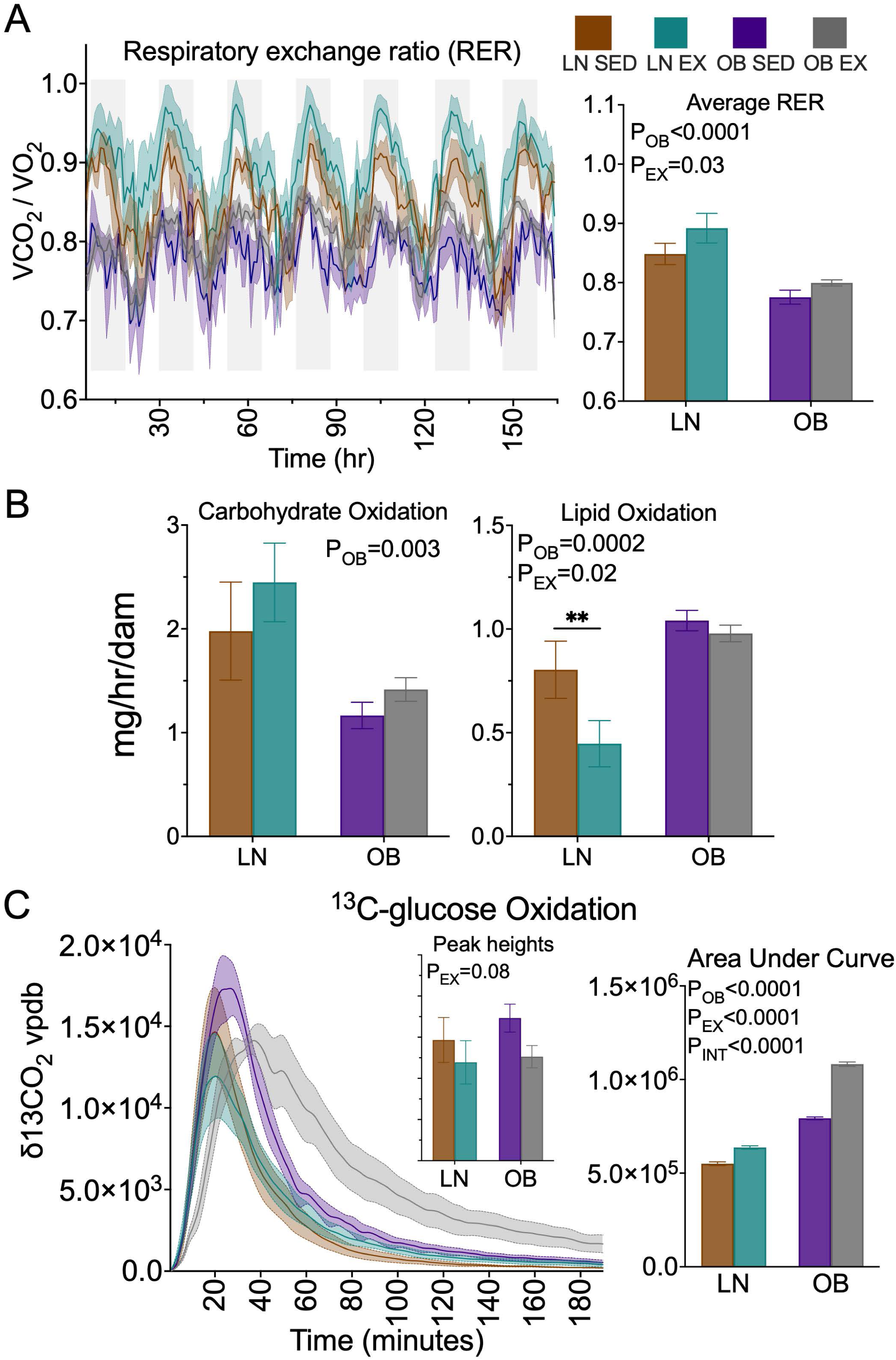
Maternal exercise affects the systemic nutrient oxidation of lactating dams differently due to maternal dietary habits. (A) Respiratory exchange ratio, a measure of how much carbon dioxide (CO_2_) the body produces relative to the oxygen (O_2_) consumed to assess the body’s fuel preference for energy, was measured in the IDC (n=5 LN EX/SED, n=6 OB SED, n=8 OB EX). (B) Carbohydrate and lipid oxidized were estimated and (C) Dams were administered a uniformly labeled ^13^C_6_-glucose (2 g/kg body weight) to quantify ^13^CO_2_ stable isotope gas exchange as a measure of whole-body glucose oxidation (n=5 LN EX/SED, n=6 OB SED, n=8 OB EX). Data are expressed as means ± SEM, and analyzed using 2-way ANOVA. Statistical significance for the main effects of obesity and exercise on the total data are denoted *P_OB_* or *P_EX_* or *P_INT_*. Asterisks represent comparison within the same group and is by *\*p<0.05, **p<0.01, ***p<0.001, ****p<0.0001*.

### 3.3 Maternal exercise remodels obesity-associated differences in mammary epithelial cell proteome

To investigate the molecular effects of maternal obesity (OB vs. LN), exercise (EX vs. SED), and their interaction (OB x EX) during lactation, we profiled isolated mammary epithelial cells (MEC) using unbiased proteomics (*Fig. 4*). Heatmap visualization illustrated broad remodeling of the MEC proteome by maternal obesity (*Fig. 4A*), with comparatively moderate exercise-related changes (*Fig. 4B*) and fewer OB x EX interactions (*Fig. 4C*). After FDR correction (adj. *p<0.05*), 945 proteins were differentially abundant between OB and LN MECs, with 604 increased and 341 decreased proteins (Supplemental File 1). EnrichR pathway analysis of the top 200 differentially aboundant proteins revealed distinct adaptations to maternal OB. MECs from OB dams displayed increased abundance of protein translation machinery (RPL, RPS, and EIF families), milk proteins (CSN2, CSN3, WAP, GLYCAM1, CSN1S1), and secretory proteins (XDH, BTN1A1). Enriched pathways included proteins for oxidative phosphorylation (ATP5 and NDUF families), fatty acid β-oxidation (ACOX3, ETFA/B), vesicle transport, and responses for endoplasmic reticulum and oxidative stress (Table 1). Maternal OB was also associated with increased lipid transport and modifying enzymes (SLC27A3, CD36, ACOT1, FADS1, LPGAT1, FPPS, GPAT1, ELOVL2/5). Conversely, decreased abundances were observed for enzymes supplying cytosolic reducing power and glycolytic flux (PFKL, G6PD1, ME1, MDH2, ALDOA/C, TKT, TALDO1, TKTL2, RPIA, LDHD), as well as the pentose-phosphate shunt (RPIA, TPI1, TALDO1, TKT), aerobic respiration (NDUF family, MDH2), mono- and dicarboxylic acid metabolism, and basal membrane integrin signaling (Laminins LAMA2, LAMB1, LAMC1). Additionally, mitochondrial complex I subunits (NDUF family) were bidirectionally regulated, suggesting remodeling or specialization of electron transport chain function in OB MECs.

**Figure 4:**
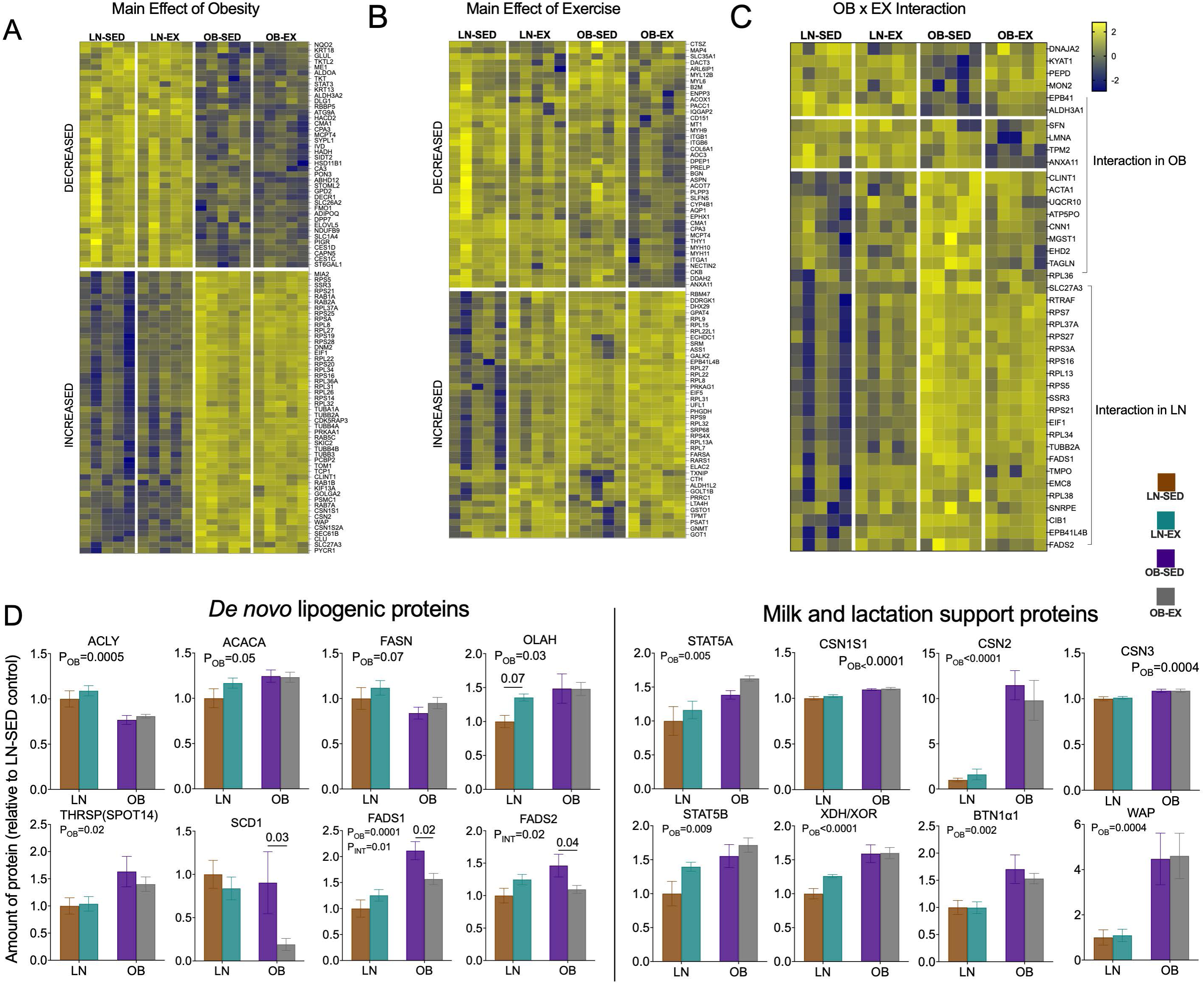
Maternal exercise restores cellular metabolic and other proteins necessary for mammary epithelial cell function. Heat maps show differentially expressed proteins (DEPs) in mammary epithelial cells due to (A) maternal obesity or (B) maternal exercise using GraphPad Prizm 10.6.0. (C) Quantification of selected proteins grouped by functional category. *De novo* lipogenic proteins (ACLY, ACACA, FASN, OLAH, SCD1, FADS1, FADS2, LPL) showed significant increases in OB dams, consistent with enhanced fatty acid synthesis and desaturation activity. Milk proteins (CSN2, WAP, LALBA, XDH/XOR, BTN1α1) were upregulated in OB dams. Regulatory proteins (THRSP/Spot14, STAT5A, and STAT5B) were elevated under maternal obesity, indicating activation of transcriptional and signaling pathways supporting lactation. Data are presented as mean ± SEM relative to LN-SED controls. Significant effects of obesity (*P_OB_*) and/or interaction with exercise (*P_INT_*) are indicated on individual graphs.

**Table 1:**
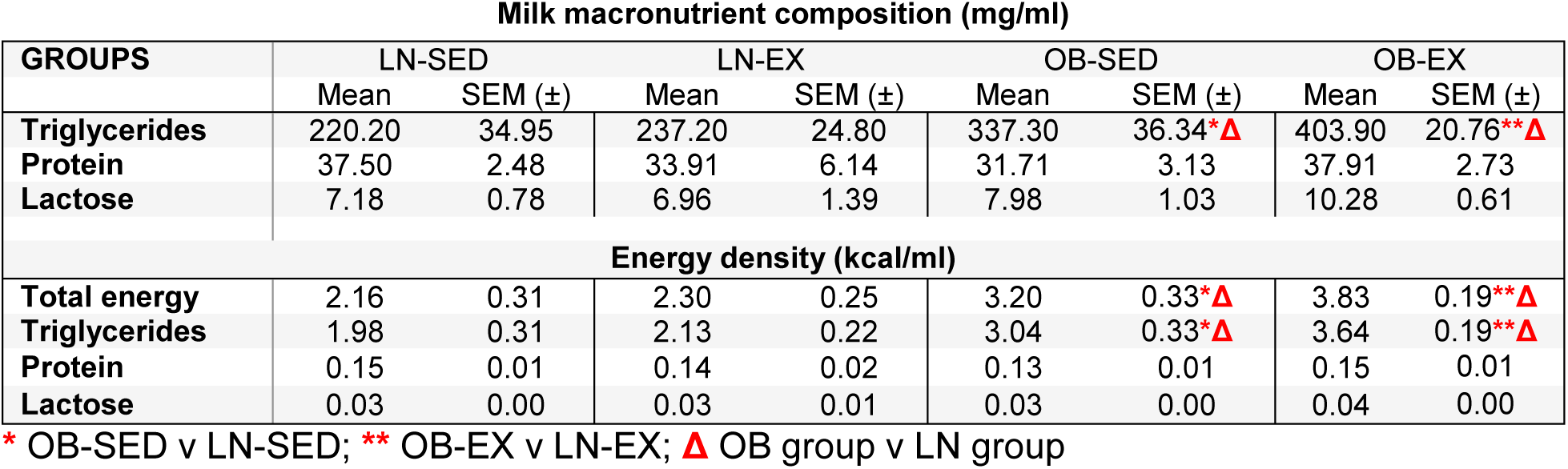
Milk macronutrient composition and calculated energy densities among groups at lactation day 12. Data presented as the mean and SEM and n=5/group, differences denoted using 2-way ANOVA with Tukey multiple testing correction.

Notably, the fatty acid biosynthesis pathway was reduced in MECs of OB dams, although the principal enzymes of the lactating mammary gland displayed a mixed pattern (*Fig. 4D*). ACLY (ATP Citrate Lyase, *P*_OB_=0.0005) was significantly reduced, while FASN (Fatty Acids Synthase, P_OB_=0.07) strongly trended lower. In contrast, ACACA (Acetyl CoA Carboxylase alpha, P_OB_=0.05), OLAH -the MEC specific enzyme that cleaves MCFA from FASN (Oleoyl-ACP hydrolase, P_OB_=0.03) (45), and THRSP that modulates the FASN rate of MCFA catalysis (Thyroid Hormone Responsive Protein Spot14, P_OB_ = 0.02) (46), were significantly increased in OB dams. STAT5A and STAT5B, transcription factors essential for lactogenesis and milk production (47), were likewise elevated in MECs from OB dams (P_OB_=0.005 and 0.009, respectively). This mixed regulation is consistent with prior reports of reduced milk medium-chain fatty acids (MCFA) in human mothers with obesity (48), suggesting that altered MEC abundance of *de novo* FA synthesis enzymes may contribute to milk fat compositional shifts observed in these mothers. Together, these findings suggest maternal OB coordinately prioritizes protein translation and FA oxidation, while suppressing glycolysis, the pentose phosphate pathway, and specific nodes of *de novo* lipogenesis.

The main effect of maternal exercise on MEC protein abundances was less pronounced than the robust adaptations observed with maternal obesity (*Fig. 4B*). No proteins passed the multiple testing correction (FDR < 0.05), likely reflecting the variance introduced by the dominating effect of maternal OB. To assess the exercise-related differences, we therefore examined proteins with unadjusted *p*<0.05, identifying 270 significantly altered proteins (158 increased, 112 decreased; Supplemental File 1). Among those increased by maternal EX, protein translation machinery (RPL, RPS, EIF families) and secretory pathways were strongly represented. In particular, proteins involved in intra-Golgi vesicle transport and COPII-coated cargo budding (RAB1A, COPA/B/G, SEC24A/D, USO1, PREB, ARCN1) were elevated, highlighting enhanced vesicle trafficking capacity required for milk secretion. Maternal EX also increased proteins supporting redox balance and metabolic reducing equivalents (NADPH/NADH regeneration via PHGDH, ALDH1L2, MDH1, ENO1, GPD1L) and nucleotide biosynthesis pathways (QNG1, PGM2/3, GMPPB, NANS, OLA1). On the other hand, proteins reduced by maternal EX were enriched for fatty acid metabolism (EPHX1, MGLL, ACOT7, ACOX1, ACADS, DECR1, SLC27A3) and regulators of cellular adhesion and migration (FGF1, ITGB1, CD151, FLNA, ITGA6, ACTN4, THY1, ENG). Together, these findings suggest that maternal exercise during lactation selectively enhances the translational and secretory machinery of MECs while promoting intracellular redox capacity needed for lipogenesis, and in parallel reduces lipid catabolism and cellular motility programs. These adaptations during exercise are consistent with lactation sufficiency, prioritizing milk synthesis and secretion over fatty acid turnover.

Analysis of the OB x EX interaction identified 71 significantly different proteins (*p*-≤0.05, unadjusted), with six increased and 65 decreased (*Fig. 4C*), too few for meaningful enrichment analysis. To better interpret these changes, we examined exercise effects within each maternal phenotype. In lean dams, maternal exercise altered 93 proteins, whereas in obese dams, exercise affected 198 proteins, indicating a stronger remodeling influence in the OB background. Within OB MECs, exercise increased proteins linked to protein translation (RPL, RPS, MRPL, and EEF families), intra-Golgi vesicle transport and COPII-coated vesicle loading and budding (COP, RAB, and SEC families, SAR1A, USO1, PREB, and TRAPPC11), glycolysis (PFKFB2, LDHA, PFKL, TPI1, GNPDA1, PKM, PKLR, PGK1, ALDOC, ENO1, PGM1), and branched-chain amino acid metabolism (BCAT2, BCKDHA and B, MCCC1 and 2, PCCA and B). In contrast, proteins decreased by EX in OB MECs were linked to electron transport chain (NDUF Complex I and ATP Synthase Complex 5 families), fatty acid oxidation (ACAD family, ECHS1, ACSL1 4 6, ACAT1, HADHA and B, CPT2, CRAT, DECR1), eicosanoid metabolism (PTGS1, PTGR1, ACOX family, EHHADH, ACAA1A), and the Kennedy phospholipid biosynthesis pathway (SGPL1, PTDSS1, CHPT1, CEPT1). Consistent with this, fatty acid desaturases that shape milk lipid profiles (SCD1, FADS1/2) were reduced in OB-EX MECs (*p*<0.02), suggesting exercise may dampen the synthesis of MUFA and LC-PUFA under conditions of maternal obesity. Together, these protein profiles suggest that exercise during lactation in obese dams reprograms MEC metabolism away from mitochondrial lipid oxidation and lipid desaturation, while enhancing glycolytic and amino acid pathways that supply biosynthetic substrates and redox equivalents. Such remodeling provides a mechanistic basis for the changes we observed in milk macronutrient and fatty acid composition.

### 3.4 Exercise increases milk fat secretion in diet-induced obese dams

We assessed whether the milk macronutrient composition and calculated caloric density varied due to maternal OB and with exercise. Milk samples were collected on Lac12 and analyzed for total triglycerides, protein, and lactose concentrations, and the nutrient density of the milk was calculated according to established methods (49). Dams in the OB group had significantly elevated milk triglyceride concentrations relative to LN group without an effect of maternal exercise (*P_OB_*=0.0003, Table 2). OB-SED dams produced significantly greater milk triglycerides than LN-SED dams (337.30 ± 36.34 mg/mL vs. 220.20 ± 34.95 mg/mL, *p*=0.01, Table 2), while OB-EX dams had more milk triglycerides than LN-EX dams (403.90 ± 20.76 mg/mL vs. 237.20 ± 24.80 mg/mL, *p*=0.001, Table 2). No significant differences in milk protein were observed among groups. OB-EX dam had significantly more milk lactose compared to LN-EX dams (*p*=0.03). We calculated caloric densities for each milk macronutrient, finding that total energy density was significantly higher in OB milk compared to LN milk (*P_OB_*=0.0003), driven by a 1.6-fold increase in energy provided by the milk triglycerides (*P_OB_*=0.0002). OB-SED dam milk had significantly higher total energy density than LN-SED dams (3.20 ± 0.33 kcal/mL vs. 2.16 ± 0.31 kcal/mL, *p*=0.02, Table 2), and OB-EX dams had more milk triglycerides than LN-EX dams 3.83 ± 0.19 kcal/mL vs. 2.30 ± 0.25 kcal/mL, *p*=0.001, Table 2). No main effect of exercise was observed. Milk composition analyses indicate that total milk energy density, likely due to the maternal HFHS diet, was mainly responsible for the observed differences in obesity.

**Table 2:**
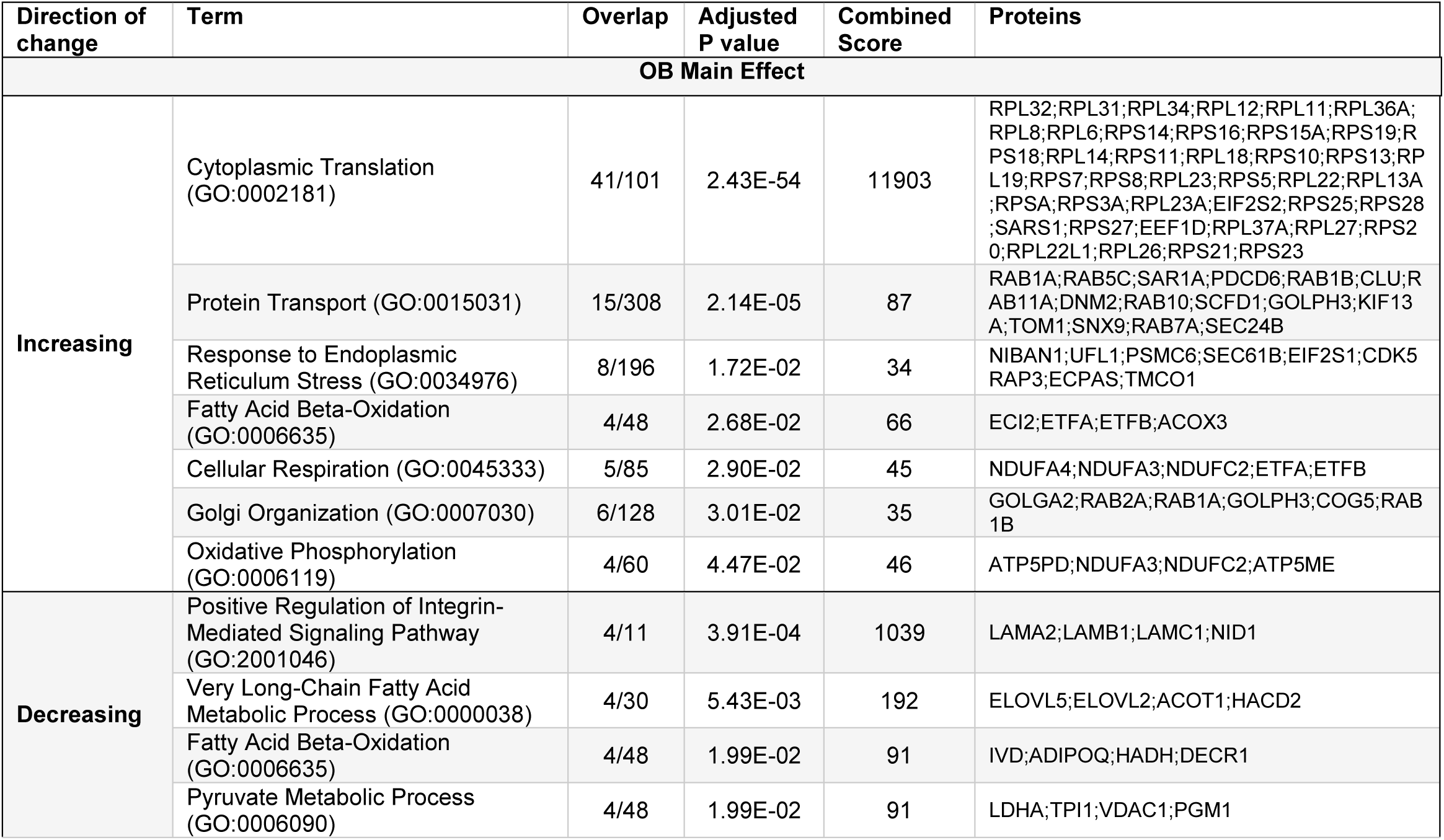
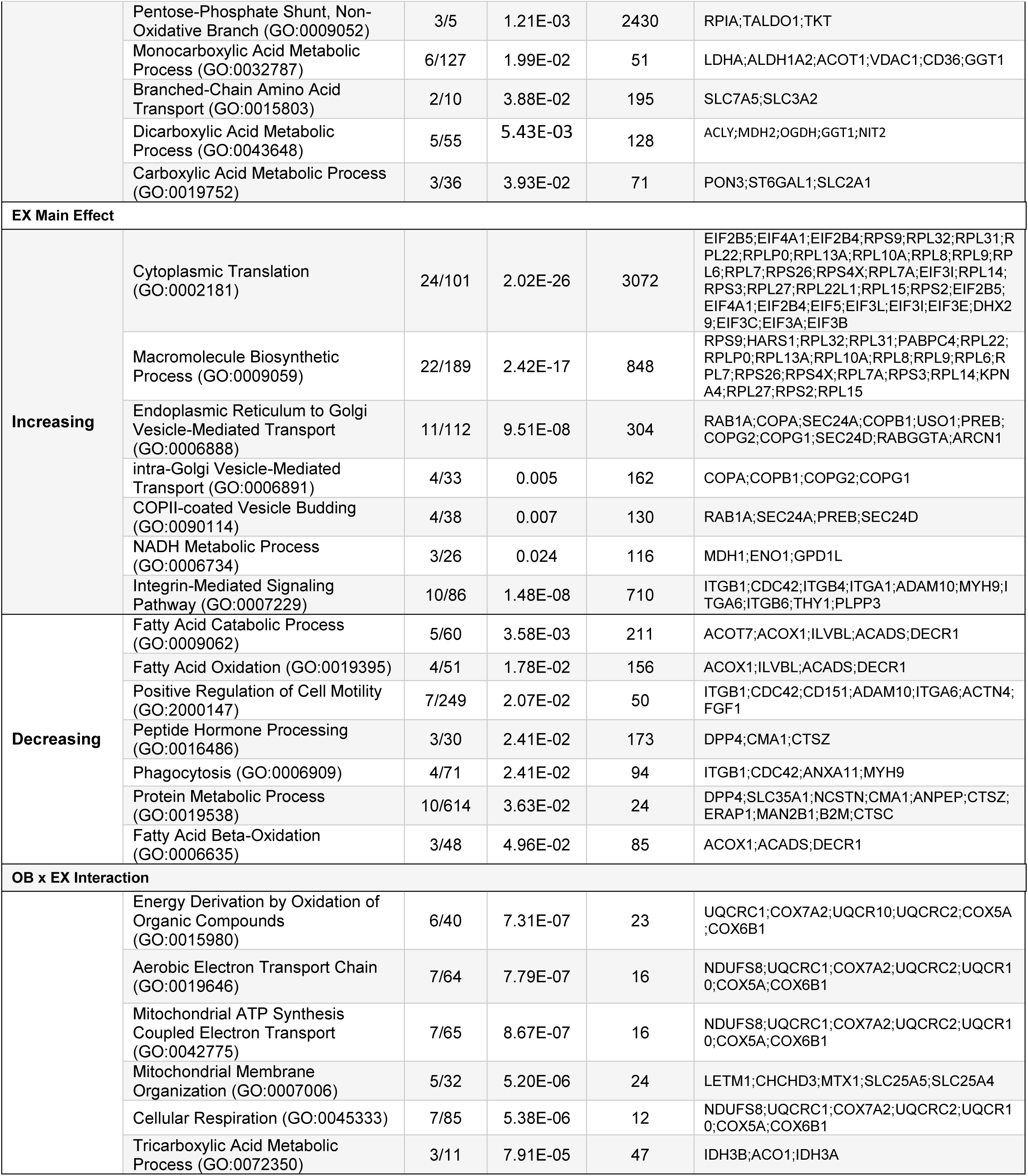
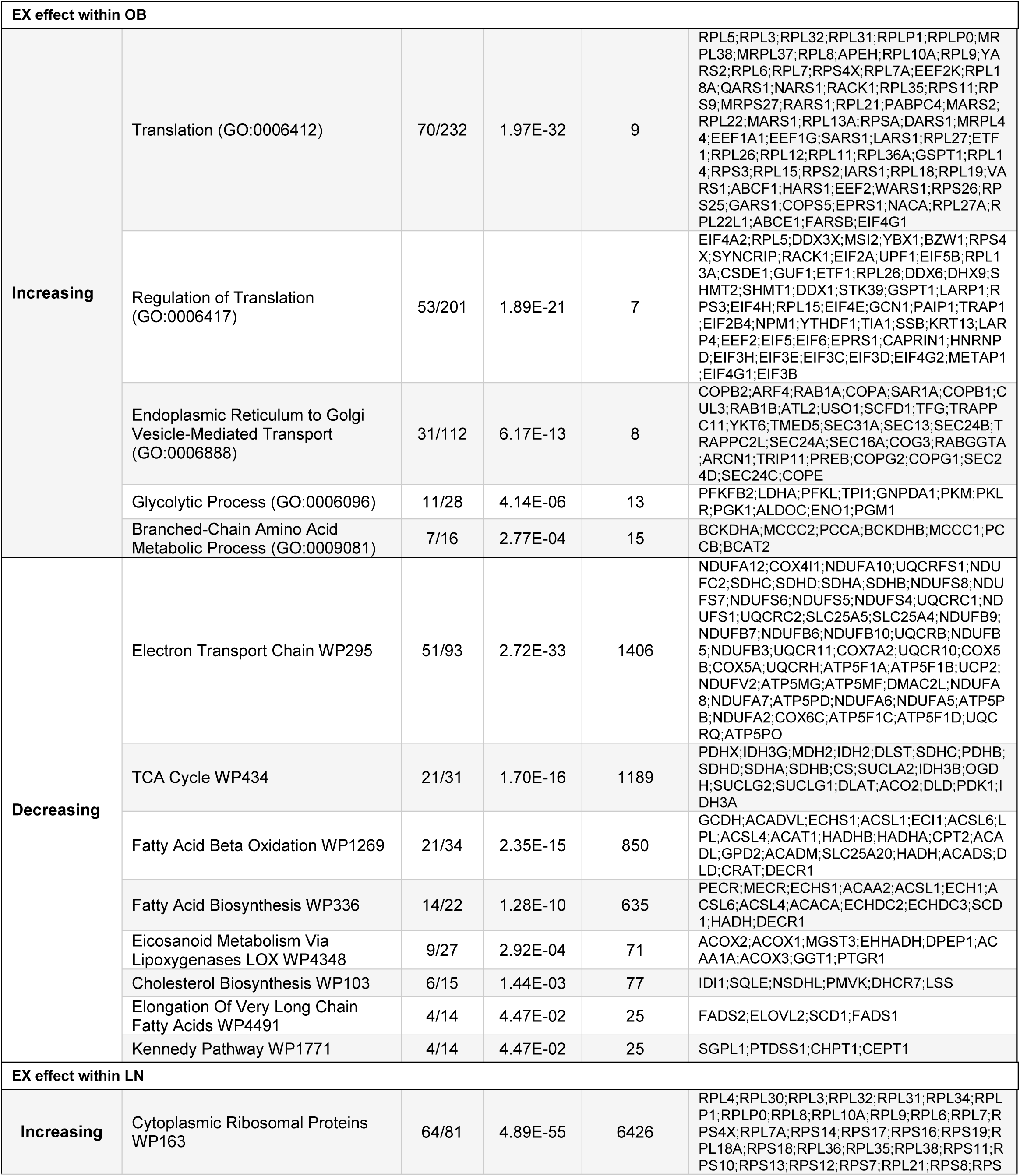
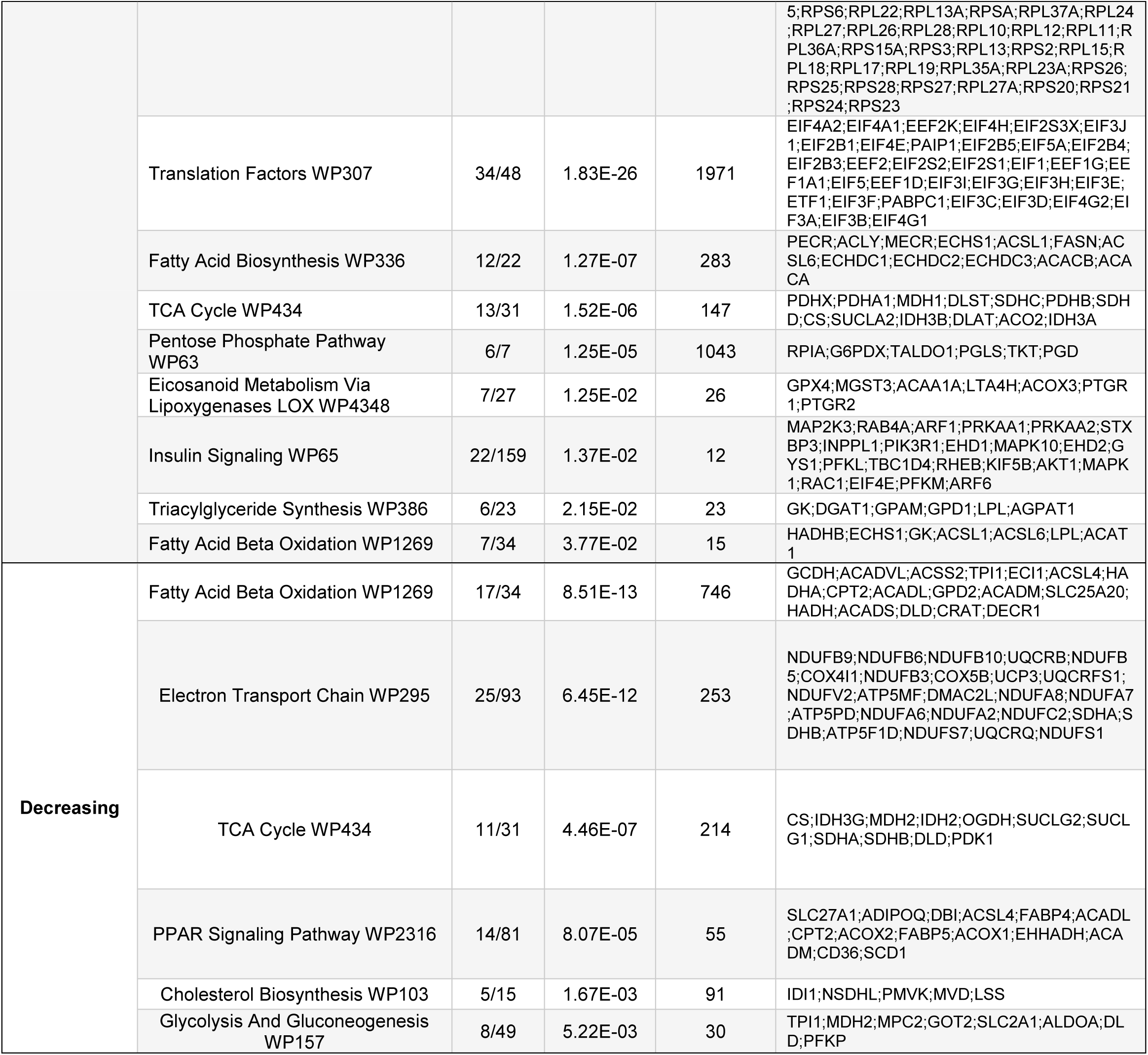
Pathways enrichment analysis performed on differentially abundant proteins from mammary epithelial cells exposed to maternal obesity and/or exercise. Only pathways with adjusted *p*<0.05 are reported.

### 3.5 Exercise during lactation boosts medium-chain fatty acids in milk and offspring circulation

We used gas chromatography mass spectrometry (GC-MS) to quantify milk fatty acid composition differences due to maternal obesity and exercise. Medium-chain fatty acids (MCFAs), derived exclusively from MEC-specific *de novo* fatty acid synthesis (46), were the most abundant fatty acid species present in Lac12 milk from OB-EX dams (*Fig. 5A*, *p*<0.0001). The sum of MCFAs was significantly increased by the main effect of obesity (*P_OB_*=0.002, Supplemental Figure S2A), particularly in OB-EX milk relative to OB-SED (*p*=0.013). The increase in MCFA in milk from OB-EX dams indicates that *de novo* FA synthesis, specifically by MECs, was enhanced by maternal exercise during obesity. Other species of FA, such as polyunsaturated fatty acids (PUFA), particularly n-6 PUFA, were significantly increased in OB-EX dams (*p*<0.01, *p*<0.05 respectively), highlighting an effect of the increase in food intake and fat loss by these dams, resulting in incorporation of preformed FA derived from the high fat diet or adipose reserves (50, 51). Analysis of the non-esterified fatty acid (NEFA) fraction in milk revealed a qualitative similarity in FA profiles for MCFA and long chain saturated FA (LC-SatFA) concentrations (Supplemental File 2).

**Figure 5:**
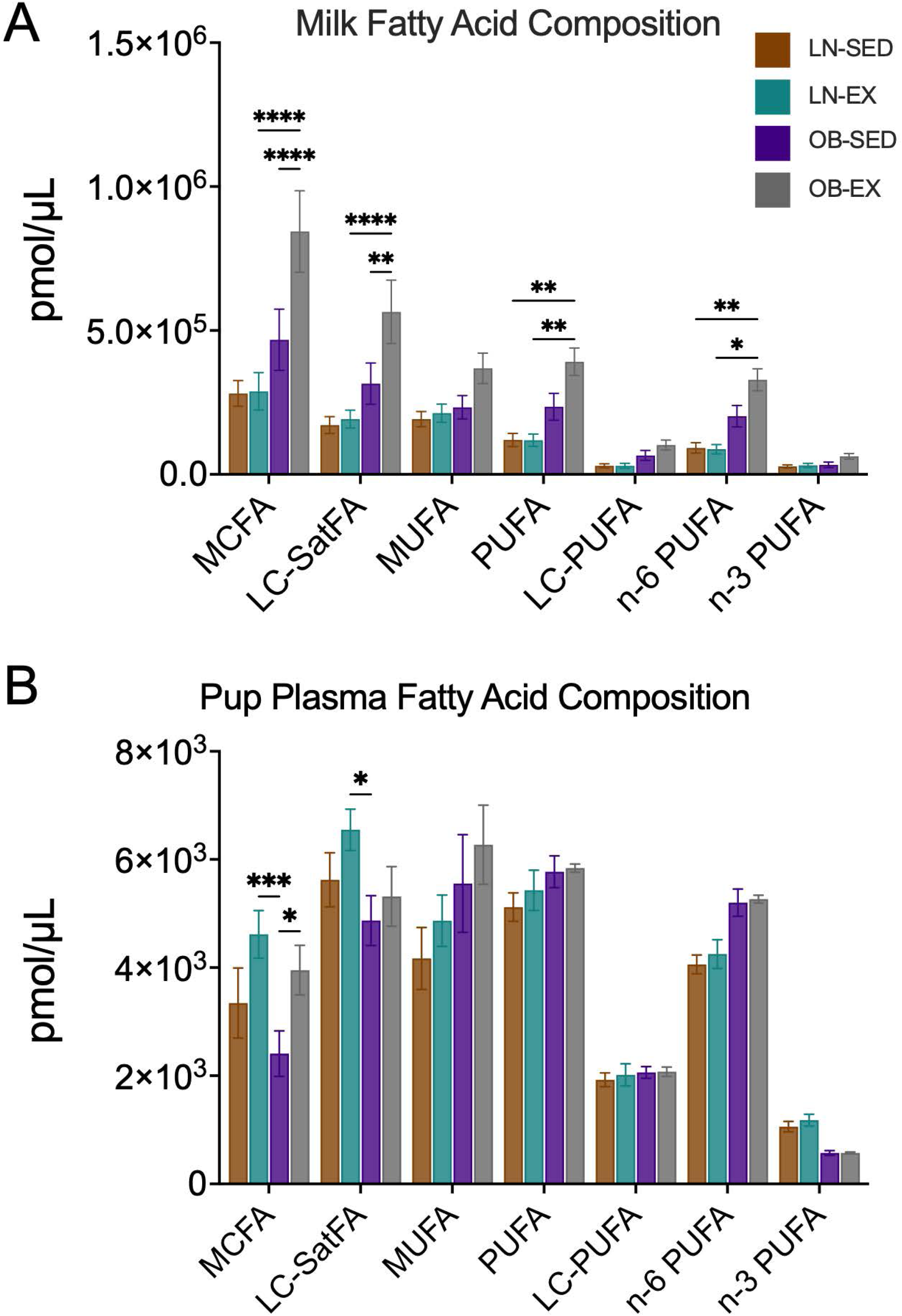
Maternal exercise increases milk medium chain fatty acids in obese dams. The effects of maternal exercise on milk fatty acid composition was analyzed using gas chromatography mass spectrometry (GC/MS). (A) Total milk fatty acids and (B) circulating plasma fatty acids were analyzed as major groups of fatty acids. Data are expressed as means ± SEM, and analyzed using 2-way ANOVA. Asterisks represent statistical significance and is denoted by *\*p<0.05, **p<0.01, ***p<0.001, ****p<0.0001*.

Given the differences in milk FA composition consumed by the litters, we investigated circulating FA profiles in pup plasma on PND12. Unlike milk, the different FA species were nearly 100-times less concentrated in pup’s plasma (*Fig. 5B*). Both LC-SatFAs and MCFAs were significantly increased in pups consuming LN-EX milk (*p*<0.001, *p*<0.05). In contrast to the concentrations in the milk, levels of MUFA and PUFA in pup’s circulation were similar in abundance with no significant differences observed among study groups. Interestingly, the quantitative sum of plasma MCFA demonstrated a main effect of maternal EX (*P_EX_*=0.02, Supplemental Figure S2B), suggesting that consumption of milk from exercising dams enhanced uptake or mobilization of these important FA energy substrates. Findings from circulating FA analyses indicate that maternal exercise enhances MCFA availability in the offspring, suggesting a potential mechanism through which maternal physical activity boosts MCFA synthesis by MECs, transmission via milk, and levels in pup circulation.

### 3.6 Post-exercise milk rescues obesity-related deficits in offspring energy expenditure and lipid oxidation

We determined the effects of maternal obesity and exercise on offspring growth, measuring body composition before and after maternal exercise intervention at PND4 and PND12 (*Fig. 6*). Prior to maternal exercise intervention, pup body weight, fat mass, and lean mass were not different between groups. By PND12, all pups had grown exponentially, as indicated by significant increases in body weight, fat, and lean mass (*P_TIME_*<0.0001). The change in weight gain over time (Δmass (g); *P_OB_*=0.0004) demonstrated a main effect of maternal obesity, while no effect of exercise was observed. Fat mass accumulation with time significantly correlated with maternal obesity, with maternal exercise not preventing fat mass expansion (*P_OB_*=0.0003). Lean mass increase with time was also primarily driven by maternal obesity, with OB pups showing significantly higher accumulation than LN pups (*P_OB_*=0.0006). Stunted offspring weight gain is often used to indicate failure to thrive (52), however, our maternal exercise regimen had no effects on overall weight gain or accumulation of body composition. Consistent with the greater caloric density of milk from OB dams (Table 2), pups consuming OB dam’s milk had greater overall growth metrics without imbalanced body composition.

**Figure 6:**
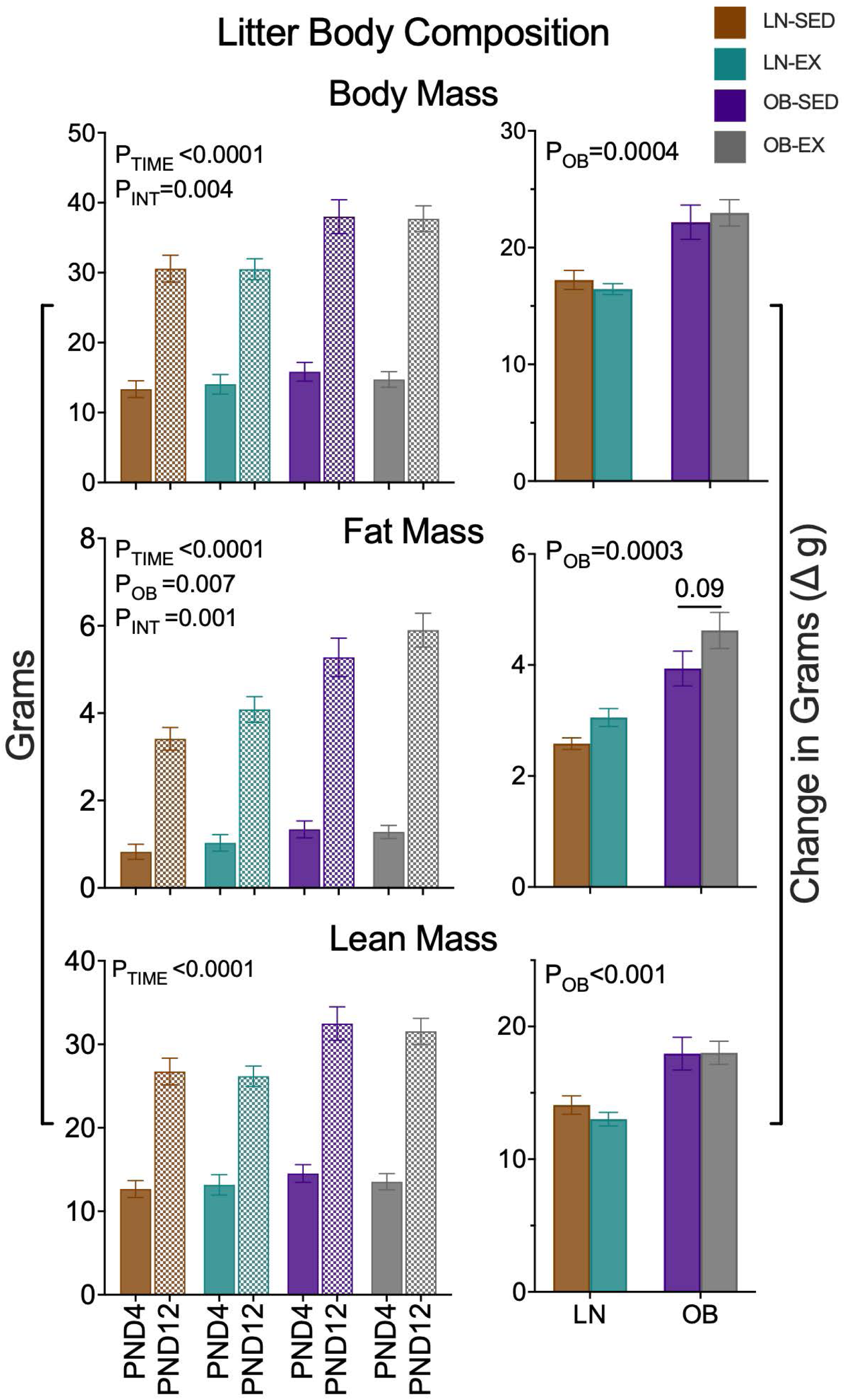
Maternal obesity increases the rate of offspring growth. Body composition of the offspring was measured to ascertain the effects of maternal exercise on offspring development and ability to thrive. Litter size was standardized to 7-8 pups per dam (n=5 LN-SED/EX, n=10 OB-SED and n=13 OB-EX). Body mass and change in body mass, fat mass and delta fat mass, and lean mass and delta lean mass were measured for each litter from all dam groups. Data are expressed as means ± SEM, and analyzed using 2-way ANOVA. Statistical significance for the main effects of obesity, exercise and interaction on the total data are denoted *P_OB_* or *P_EX_* or *P_INT_*.

Given the impact of maternal obesity and exercise on milk FA composition, we evaluated litter metabolic phenotypes using indirect calorimetry and stable isotope tracer analysis as previously described (24). Pups consuming OB milk exhibited significantly diminished energy expenditure (EE) than pups consuming LN milk (*Fig 7A*, *P_OB_*=0.012) potentially due to their larger size. However, pups consuming OB-EX milk tended to have increased total EE relative to the OB-SED group (*p*=0.09), even though these pups tended to have more fat mass (*Fig. 7*, *p*=0.09, fat mass), suggesting an improvement in promoting EE with maternal exercise. The RER measurements were less than 0.7 for all pup groups, indicated oxidation of lipids as a metabolic fuel. The main effect of maternal OB on RER was a significant increase in pups reared by OB relative to LN dams (*Fig. 78B*, *P_OB_*=0.039), indicating a significant shift in fuel utilization away from lipids. From the gas exchange data, we calculated lipid and carbohydrate oxidation rates and found the main effect of maternal obesity strongly tended to decrease lipid oxidation rates but not carbohydrates (*Fig. 7C*, *P_OB_*=0.078). Interestingly, the slight improvement in OB-EX pups suggested a possible recovery in lipid oxidation. To further quantify *in vivo* lipid oxidation, we administered an independent cohort of pups with 100mg/kg of uniformly ^13^C- labeled palmitate to quantify FA oxidation (FAO) as indicated by ^13^CO_2_ emission in the breath of pups (full trace, *Fig. 7D*). Notably, FAO was significantly increased as a main effect of maternal EX with a significant interaction driven by the LN group (*P_EX_* and *P_INT_*<0.0001), although FAO was significantly decreased overall in pups reared by OB dams (*P_OB_*<0.0001). Consistent with the slight increase in calculated lipid oxidation rate of pups consuming OB-EX milk, ^13^C- palmitate FAO was significantly increased in OB-EX pups relative to OB-SED group (*p*=0.002). This finding indicates that offspring consuming milk from maternal exercise dams, particularly OB-EX dams, had improvements to whole-body energy metabolism, energy expenditure, and lipid fuel utilization.

**Figure 7:**
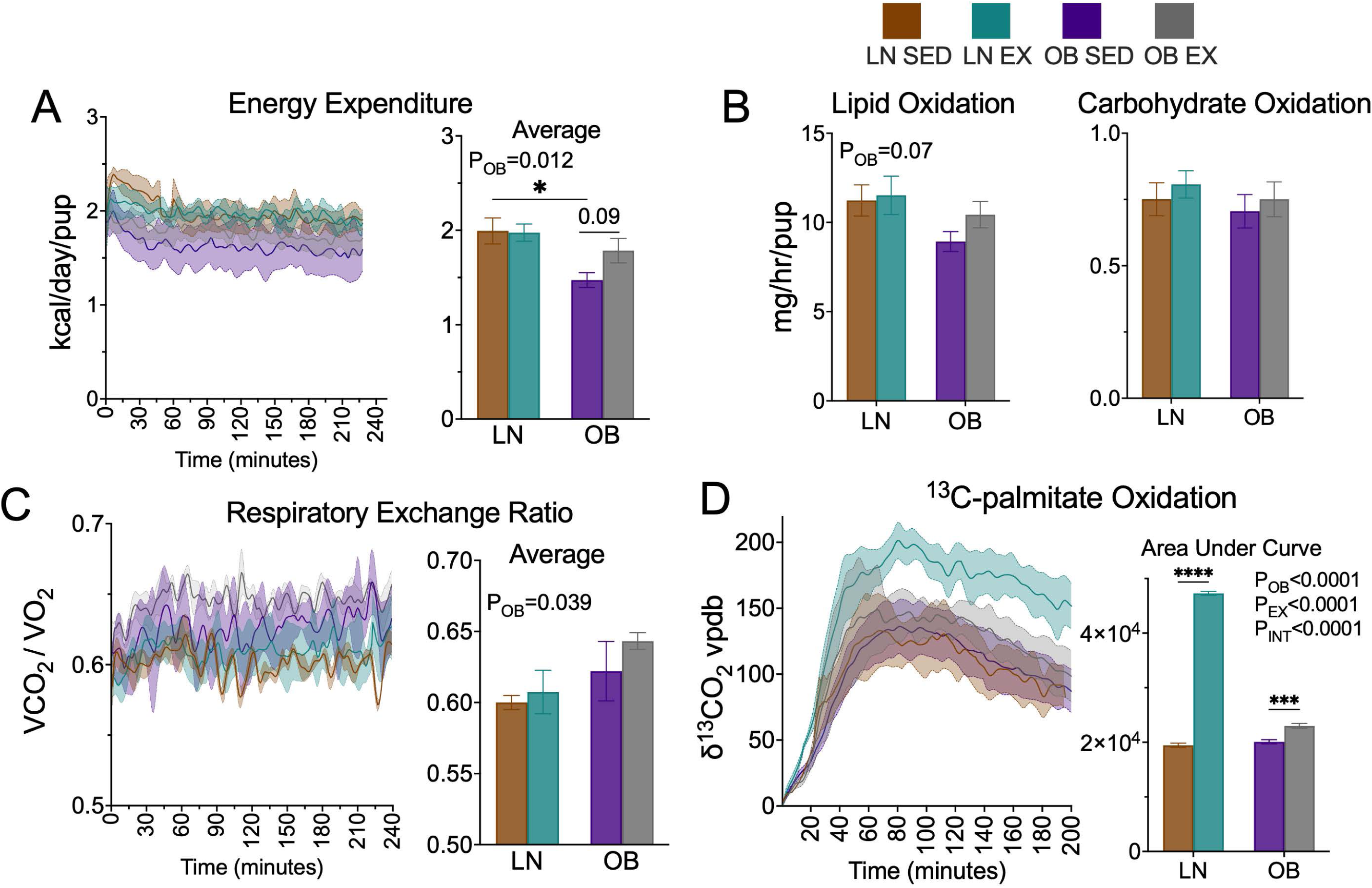
Exposure to exercised milk restores fatty oxidation and energy expenditure in OB offspring. Dams were separated from pups in the calorimetry cages within the environmental cabinet maintained at 25 °C, for about 3 – 4 h to measure litter’s (A) energy expenditure, (B) respiratory exchange ratio, and (C) lipid and carbohydrate oxidation. (D) Pups were individually administered uniformly labeled ^13^C_16_-palmitate (100 mg/kg body weight) to quantify ^13^CO_2_ stable isotope respiration for systemic nutrient oxidation (n=4 litters in LN-EX/SED, OB-EX, n=5 litters in OB-SED). Data are expressed as means ± SEM, and analyzed using 2-way ANOVA. Statistical significance for the main effects of obesity, exercise and interaction on the total data are denoted P_OB_ or P_EX_ or P_INT_. Asterisks represent statistical significance within groups and is denoted by *\*p<0.05, **p<0.01, ***p<0.001, ****p<0.0001*.

### 4.0 Discussion

This study provides new insights into how maternal exercise during lactation influences maternal physiology, MEC proteome, milk fatty acid composition, and neonatal growth and metabolism in the context of maternal obesity. While successful breastfeeding is optimal for providing the macro- and micronutrients for infant development, maternal challenges such as obesity can affect milk production and composition, impacting infant health (17, 31, 53).

Consistent with our prior work demonstrating that human maternal exercise during lactation modifies the milk metabolome and lipidome with consequences for infant body fat accumulation (32, 33), we developed a mouse model to test the physiological and molecular adaptations of the dams and their MECs to obesity and exercise. Lactation presents a high metabolic demand for the mother, just as postnatal growth is metabolically demanding for the infant. Maternal energy intake (EI) and stored nutrient mobilization offset the increased energy demands of milk production in humans and rodents (49, 54, 55). In our mouse model, the indirect calorimetry data demonstrate distinct adaptive strategies to the added cost of daily, moderate exercise, wherein the LN-EX dams lowered their energy expenditure (EE) without increasing EI, and OB-EX dams increased EI while maintaining EE reaching a higher energy balance (*Fig. 2*). This finding was consistent with mouse studies demonstrating that high fat-fed OB lactating dams had significantly greater EI and reached a higher energy balance than LN controls (49).

OB-EX dams simultaneously increased their EI and decreased body fat mass, without modifying their EE, while the OB-SED dams significantly increased their body mass driven by gains in lean mass without any decrease in fat mass (*Fig. 1* and *2*). Importantly, this adaptive response in OB-EX dams of increasing intake and mobilization of nutrient energy countered the secondary energy demand placed on the lactating dams by daily, moderate exercise. Stable isotope gas analysis studies further supported these adaptations, in that ^13^C-glucose oxidation was sustained in the OB-EX dams, potentially marking a shift from lipid fuel utilization (*Fig. 3*). This adaptation was noted in the whole milk macronutrient composition, in which the OB-EX dams produced milk with the greatest energy density, due to enhanced milk triglyceride content (Table 1). EX-mediated milk production/secretion is further supported by the consistent linear growth observed among all study pups, regardless of the exercise intervention, indicating the maternal exercise regimen preserved the offspring capacity to thrive despite the added energetic demands of exercise beyond that of lactation (*Fig. 6*). At this time, we did not evaluate maternal insulin sensitivity or lowered inflammatory status with exercise, which may exert systemic effects shown to benefit milk biosynthesis and nutrient delivery (56). Improvements via exercise in mammary gland sensitivity to systemic metabolic hormones is the subject of future studies.

Preceding lactation, MECs undergo substantial functional differentiation to accommodate copious milk synthesis and secretion under the stimulation of multiple regulatory mechanisms (4, 47, 57). Detailed proteomic analyses of milk fat globules, and immortalized mouse MECs highlight the large number of the lipid metabolic pathways, glycolytic enzymes, mitochondrial biogenesis and complex proteins, secretory machinery, and functional differentiation regulators that coordinate the complexity of milk production (58–61). Our findings suggest that maternal obesity did not, *per se*, adversely impact the differentiation of the MECs or impair their lactational capacity. This was evidenced by the significant expression of key milk proteins, including β-casein and WAP in MECs from obese dams, indicating that functional differentiation and secretory activation was preserved, despite the diet induced obesity beginning at 8-weeks of age (*Fig. 1A*). These results highlight that, rather than limiting lactational globally, molecular adaptations to maternal obesity operates through more complex mechanisms, implying that exercise may enhance the capacity of MECs in support of milk production and secretion.

At the proteomics level, maternal obesity had a dominant effect on MEC protein profiles, particularly in pathways not traditionally associated with lactation, such as fatty acid oxidation and oxidative phosphorylation via mitochondrial complex I and V components. We found exercise reduced proteins involved in lipid oxidation, fatty acid desaturation, and phospholipid biosynthesis, particularly in MECs isolated from OB dams. At the same time, maternal exercise enhanced translation machinery and vesicle budding pathways, processes that are essential for milk protein synthesis and overall secretory capacity. This exercise-mediated shift suggests a metabolic remodeling away from mitochondrial lipid catabolism towards glycolytic and amino acid derived biosynthetic precursors. To that end, an increased representation of glycolytic and branched-chain amino acid metabolism pathways may provide both biosynthetic substrates and reducing equivalents (NADPH/NADH) to support *de novo* lipogenesis and protein glycosylation, processes that are central to both macronutrient and bioactive milk contents. Metabolic pathways such as the pentose phosphate pathway, ETC and oxidative phosphorylation are responsible for generating the energy needed for the function of the cells (62). Through Metscape analysis, Jaswal *et al*, 2020, showed similar pathways in the buffalo MEC line (58), revealing similar pathways in mammary cells of different animal models. Enrichment analysis of bovine MECs infected by bacterium *S*. *agalactiae* MEC showed decreases of glycolysis and gluconeogenesis, pentose phosphate pathways and pyruvate metabolism, suggesting that inhibition of these pathways diminished ATP production, a key intracellular currency needed for lipogenesis and milk protein synthesis (63). Moreover, the observed decreases in fatty acid desaturases (SCD1, FADS1/2) with maternal exercise under obese conditions suggests a potential mechanism for altered milk fatty acid composition. By establishing a mechanistic framework in a controlled mouse model, these data provide a translational bridge to our prior and ongoing studies of human exercise induced changes to the breastmilk lipidome and thermogenic metabolites (33).

Milk fatty acid (FA) composition reflects contributions from three sources: *de novo* synthesis in MECs, mobilization of maternal adipose stores, and direct dietary FA incorporation (64–66). DNL encompasses several lipid biosynthetic arms, including cholesterol biosynthesis, phospholipid and triglyceride synthesis (67). Within this broader program, *de novo* fatty acid synthesis refers specifically to the cytosolic pathway in which glucose-derived citrate is converted to acetyl-CoA by ACLY, carboxylated to malonyl-CoA by ACACA, and acyl chain elongation occurs in the presence of malonyl-CoA and NADPH by FASN, typically producing palmitate as the product (68, 69). The lactating mammary gland is a unique setting in which this pathway is highly active, as palmitate and MCFA, through the actions of OLAH and THRSP of FASN catalysis (70), provide major energy substrates for neonatal lipid oxidation. In our study, proteomics analyses revealed that maternal obesity reduced key enzymes of *de novo* FA synthesis (ACLY, FASN), consistent with prior evidence that obesity shifts MEC lipid metabolism away from glucose-derived carbon and toward dietary lipid uptake (19, 71). At the same time, OLAH, the enzyme that enhances MCFA release from FASN, and THRSP which enhances FASN catalysis, were increased in MECs of OB dams. These findings suggest selective regulation within the pathway during lactation. Importantly, maternal exercise during lactation partially restored abundance of selected lipogenic proteins and enhanced glycolytic and amino acid pathways that supply biosynthetic precursors and reducing equivalents (NADPH) for FA synthesis (Table 1).

Glucose is considered a primary substrate for *de novo* fatty acid synthesis (72). Using ^13^C-glucose tracer oxidation assay we observed sustained oxidation in the OB-EX dams, rather than a rapid oxidation and quicker ^13^CO_2_ decay rate relative to the OB-SED dams (Fig 3C). We speculate that this sustained oxidation might reflect increased entry of glucose carbon into the ACLY - ACACA - FASN pathway for *de novo* FA synthesis, thereby supporting greater milk MCFA output. Consistent with this, milk from OB-EX dams contained significantly higher levels of MCFA, a pattern that was observed in both the total FA and NEFA fractions (Supplemental file 2). More detailed studies are needed to address this possibility, however, at 24-hours post administration of the ^13^C-glucose tracer, negligible ^13^C enrichment was observed in the milk FA composition, suggesting that the tracer was secreted into the milk and was therefore washed out (data not shown).

Previous work by Harris, et al., showed that maternal exercise in humans alters the milk metabolome, particularly pathways related to lipid metabolism, further substantiating the findings observed here in the murine model (73). A recent review by Moholdt and Stanford, 2024, has also highlighted the various metabolic benefits and obese prevention capabilities of breastmilk and breastfeeding (74). Another notable outcome of this study is the partial correction of the n6:n3 fatty acid ratio in pups from OB-EX dams (Table 2). Elevated n6:n3 ratios are associated with heightened inflammatory risk and are a known feature of obesity-induced metabolic dysfunction (75). While maternal high fat feeding significantly increased this ratio in both total and non-esterified fatty acid pools in pup plasma, exercise attenuated this imbalance. These findings suggest that maternal exercise may reduce the pro-inflammatory lipid exposure in neonates born to obese mothers, providing a nutritional environment more conducive to healthy metabolic development.

Broad differences in the proteomics pathways for protein translation components (RPL, RPS, EIF families) were observed in the isolated MECs (*Fig. 4*). Interestingly, gene expression patterns did not fully align with the levels of selected proteins measured by proteomics (Supplemtental Figure S3). This finding indicates another level of complex regulation is in place, potentially regulation of mRNA loading into the ribosomes by micro-RNAs. Elegant work by Heinz et al., demonstrated that constitutive overexpression of micro-RNA mir-150-5p significantly decreased protein levels of ACACA, FASN, and OLAH by suppressing transcript levels, resulting in significantly lower levels of MCFA in the milk (76). Notably, maternal exercise appeared to enhance or restore the expression of key milk protein genes, including Csn2, Wap, and Lalba, while no differences in their protein levels were observed. At present time, we are uncertain whether the EX-mediated increase in OB MECs for *de novo* FA synthesis genes (Srebf1c, Olah, Thrsp) represents potential increased sensitivity to circulating hormones or nutrient signals. Regardless, these findings suggest that maternal exercise during lactation, particularly in OB dams, may enhance the functional capacity of the mammary epithelium to support milk quality.

Maternal obesity/high fat feeding significantly altered pup body composition, with litters born to OB dams exhibiting greater fat and lean masses relative to lean controls. Despite all offspring beginning with similar total body weights, pups born to OB dams exhibited a significantly greater fat mass at PND4 without changes in lean mass. This finding is consistent with large for gestational age babies frequently coincident with maternal overweight and obesity in humans (31, 77, 78). Interestingly, maternal exercise in the OB-EX litters tended to have increased fat mass, likely due to the increase caloric density driven by greater milk triglycerides. While increased infant adipose accumulation is associated with elevated risk for developing obesity later in life (79–81), human studies of infant systemic metabolism with maternal interventions like exercise are limited. Stunted offspring weight gain is often used to indicate failure to thrive (52), however, our maternal exercise regimen had no effects on overall weight gain or accumulation of body composition. Consistent with the greater caloric density of milk from OB dams (Table 2), pups consuming OB dam’s milk had greater growth metrics without imbalanced of body composition. Beyond body composition, we show that maternal obesity impaired systemic metabolism in pups, evidenced by reduced energy expenditure, decreased lipid oxidation, and a significantly increased RER. Neonatal reliance on FAO is a well-established survival mechanism, as both ketogenesis and gluconeogenesis are essential for maintaining glucose homeostasis in the first two weeks of life; disruption of either pathway is uniformly lethal (82–84). In line with this, Bairam et al. 2012, reported remarkably low RER values in neonatal rats, averaging ∼0.46 at PND4 and increasing only to ∼0.70 by PND12, consistent with a dominant reliance on lipid oxidation (85). We have recently reported similar RER observations in neonatal mice (24), reinforcing the physiological basis of these low values. This dependence on fat-derived substrates supports systemic energy balance, intestinal maturation, and postnatal organ development, including cardiomyocyte mitochondrial remodeling driven by ketone bodies (86, 87). Such metabolic programming ensures neonatal survival when maternal nutrient delivery is limited and aligns with recent evidence that maternal-fetal communication directs these adaptations (88). Thus, our observed low RER values likely capture a necessary and conserved feature of neonatal metabolism rather than an experimental artifact. In our mouse model, maternal exercise during lactation tended to increase the pup EE through significantly greater lipid oxidation measured by ^13^C-palmitate tracing. Although these adaptations are consistent with improved metabolic flexibility in adults (89), in the context of postnatal anabolic growth and development, systemic metabolism of 12-day old pups was responsive to macro- and micronutrient changes in milk composition due to maternal exercise.

In humans, maternal resistance exercise during pregnancy increased infant resting energy expenditure by >35% versus aerobic exercise or no exercise at 1 month (56). In our recent clinical studies, maternal physical activity during lactation associated with reductions in milk metabolites linked to increased infant obesity risk (32), while greater levels of milk-derived signaling lipids associated decreased risk (33). Similarly, findings in mice demonstrate that maternal perinatal exercise shapes pup energy metabolism, reducing susceptibility to obesity and insulin resistance in later life (38, 39, 90). Li et al., reported that exercising dams during gestation increased secretion of the adipokine SERPINA3C, which improved offspring glucose tolerance and insulin sensitivity via KLF4-mediated suppression of adipose inflammation (91). Moreover, other rodent studies demonstrate that maternal exercise induces circulating “exerkines” such as Apelin, which enhance fetal and neonatal brown adipose tissue development and protect against high fat diet-induced metabolic dysfunction (92). Although many of these studies included gestational exercise, our findings highlight that lactation alone represents a critical and modifiable window where maternal behavior influences offspring metabolic programming.

Several limitations of this study should be acknowledged. First, while indirect calorimetry captured maternal energy expenditure during lactation, exercise-associated energy costs could not be measured because dams were removed from metabolic cages during treadmill bouts. This likely led to an underestimation of total maternal energy expenditure. Second, proteomic analyses were performed for total protein levels without assessing post-translational modifications, like phosphorylation, which are often critical to metabolic enzyme activation status. The high-fat high-sucrose diet enriched in linoleic acid from corn oil effectively induced obesity but does not capture the heterogeneity of human maternal diets, potentially limiting direct translation. While ^13^C-glucose tracer studies provided novel insight into systemic substrate oxidation in dams and pups, enrichment of labeled carbon into milk fatty acids was not quantifiable, preventing direct assessment of maternal glucose-to-MCFA flux into milk. More broadly, species differences in milk composition, including a greater representation of medium-chain fatty acids in mice compared to humans, places limitations on translational interpretation. Offspring phenotyping was restricted to postnatal day 12, which represents a critical anabolic window, but longer-term follow-up is required to determine whether the metabolic adaptations observed persist beyond the suckling period. While the forced treadmill protocol ensured consistency in exercise dose, it differs from voluntary physical activity and may introduce stress responses that could influence maternal physiology. Although significant differences were identified, certain measurements may have been underpowered, limiting the ability to detect additional effects. Together, these limitations underscore the need for longitudinal studies across lactation, more refined metabolic tracing approaches, and human trials to fully define how maternal exercise interacts with obesity to shape milk composition and offspring metabolic outcomes.

In conclusion, this study integrates maternal physiology, MEC proteomics, milk fatty acid composition, and offspring metabolic phenotypes, providing a mechanistic view of how maternal obesity and exercise interact during lactation. Maternal obesity imposed broad remodeling of MEC protein networks, reducing enzymes critical for *de novo* fatty acid synthesis while increasing reliance on dietary lipid uptake. These molecular changes corresponded with altered milk fatty acid profiles, characterized by higher triglyceride content and a pro-inflammatory n6- to n3-FA balance, which were reflected in pup plasma. Offspring nursed by obese dams exhibited greater fat mass and impaired energy metabolism, consistent with early-life programming of adiposity. Importantly, maternal exercise during lactation partially mitigated these effects by enhancing MEC translational and vesicle transport capacity, restoring elements of glycolytic and lipogenic metabolism, shifting milk fatty acid composition toward more oxidizable substrates, and reprogramming offspring energy expenditure and fatty acid oxidation. Cumulatively, these findings highlight lactation as an intervention window in which maternal activity and diet could be manipulated to influence both milk composition and neonatal metabolism to optimize the health trajectory of the mother-infant dyad.

## Acknowledgements

MCR supported by R01HD117197, R56DK139005-01A1, the Oklahoma Center for Adult Stem Cell Research, the Presbyterian Health Foundation Bridge Fund and Team Science Awards, and the College of Medicine Alumni Association. MK supported by R24GM137786 (IDeA National Resource for Quantitative Proteomics) and P20GM103447 (Oklahoma INBRE).

## Authors contributions

MCR, EMI, DAF, KS, GKD conceived and designed research; GKD, RV, MCR analyzed the data, GKD, RV, SD, AM, KH, JWF, MK, GM performed the experiments; GKD, RV, MCR interpreted results of experiments; GKD, MCR prepared figures; GKD, MCR drafted manuscript; GM, RV, KH, CMM, EMI, DF, KS, DAF, MCR edited and revised the manuscript; MCR, DAF, EMI approved final version of the manuscript.

---

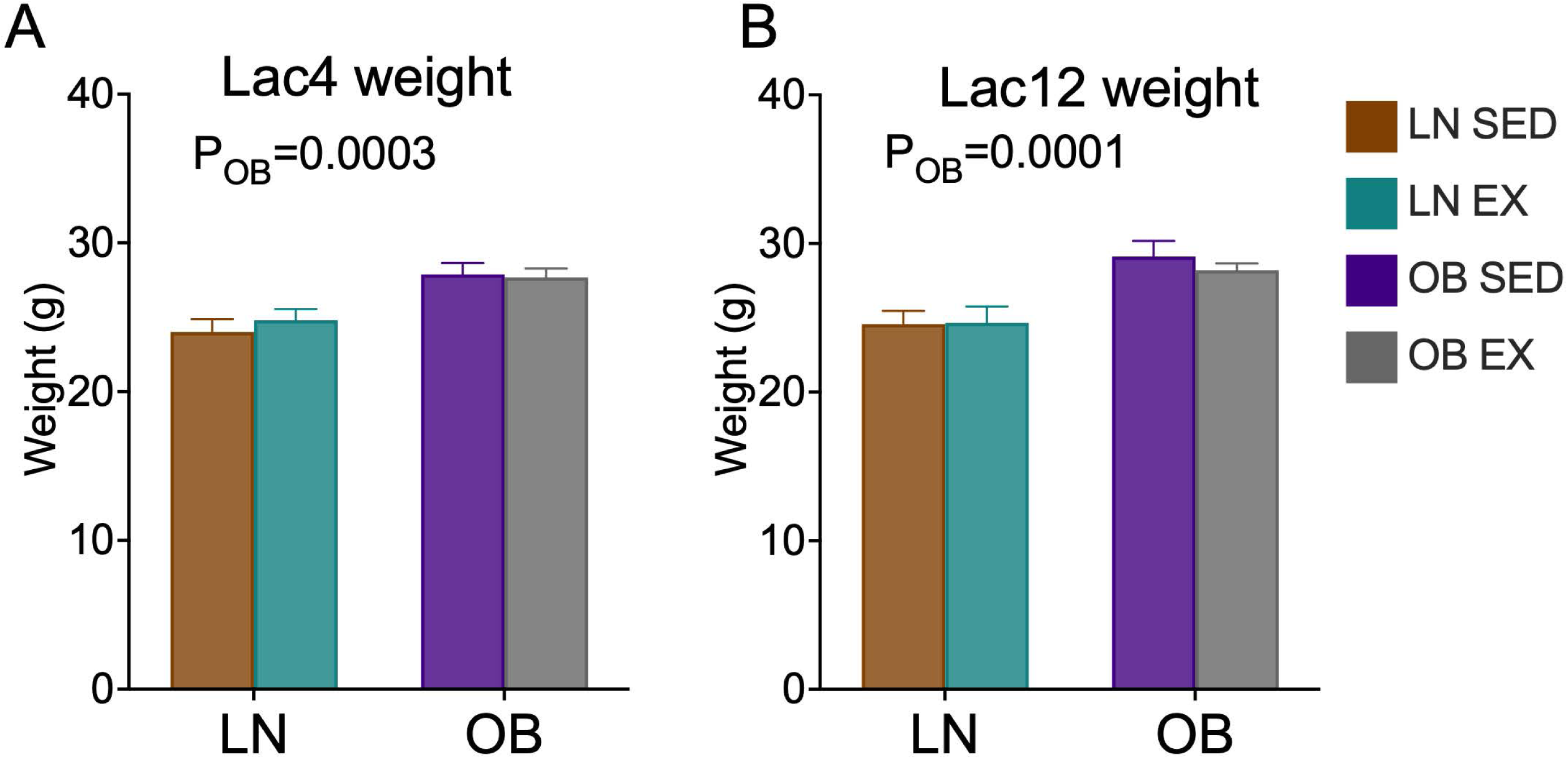

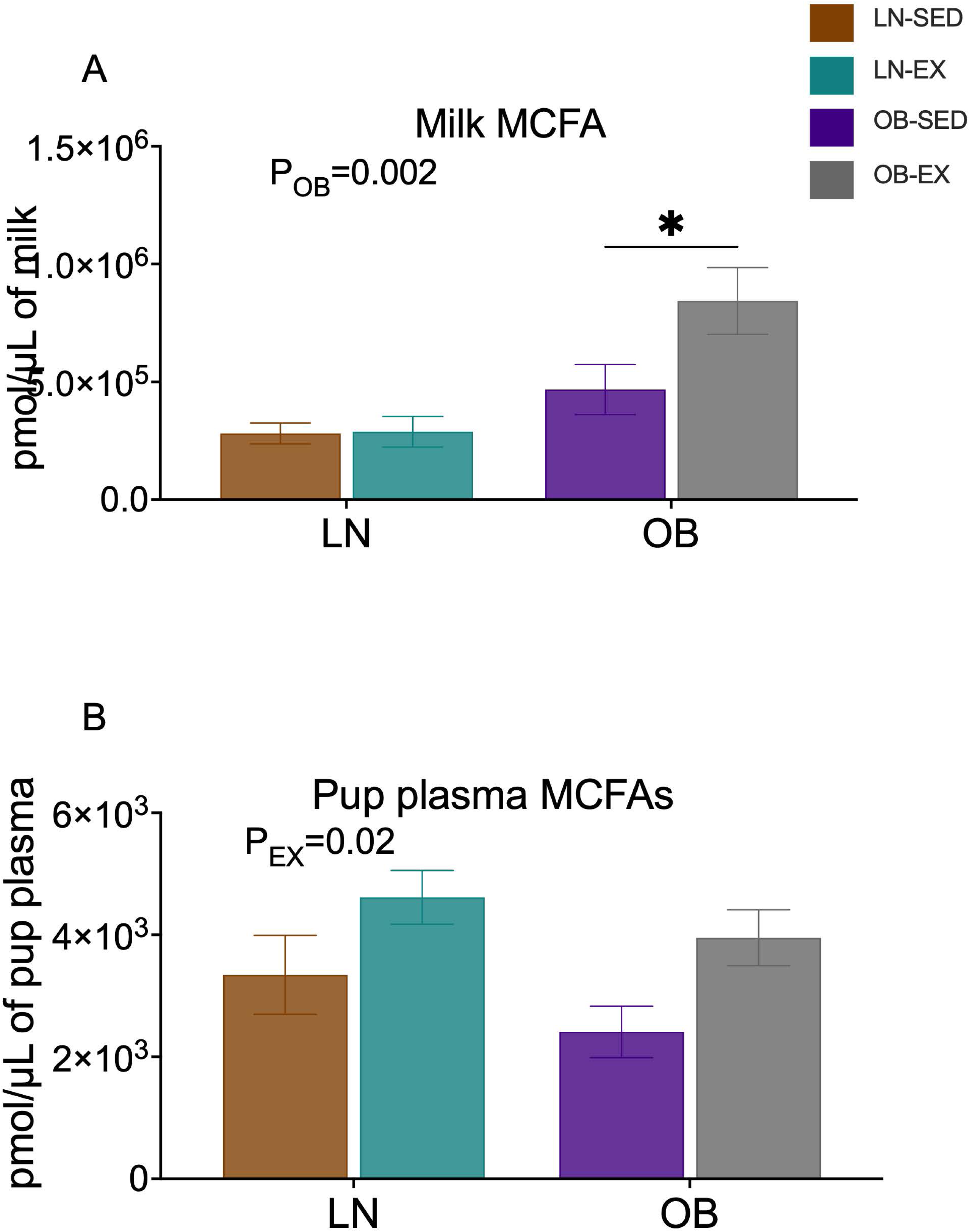

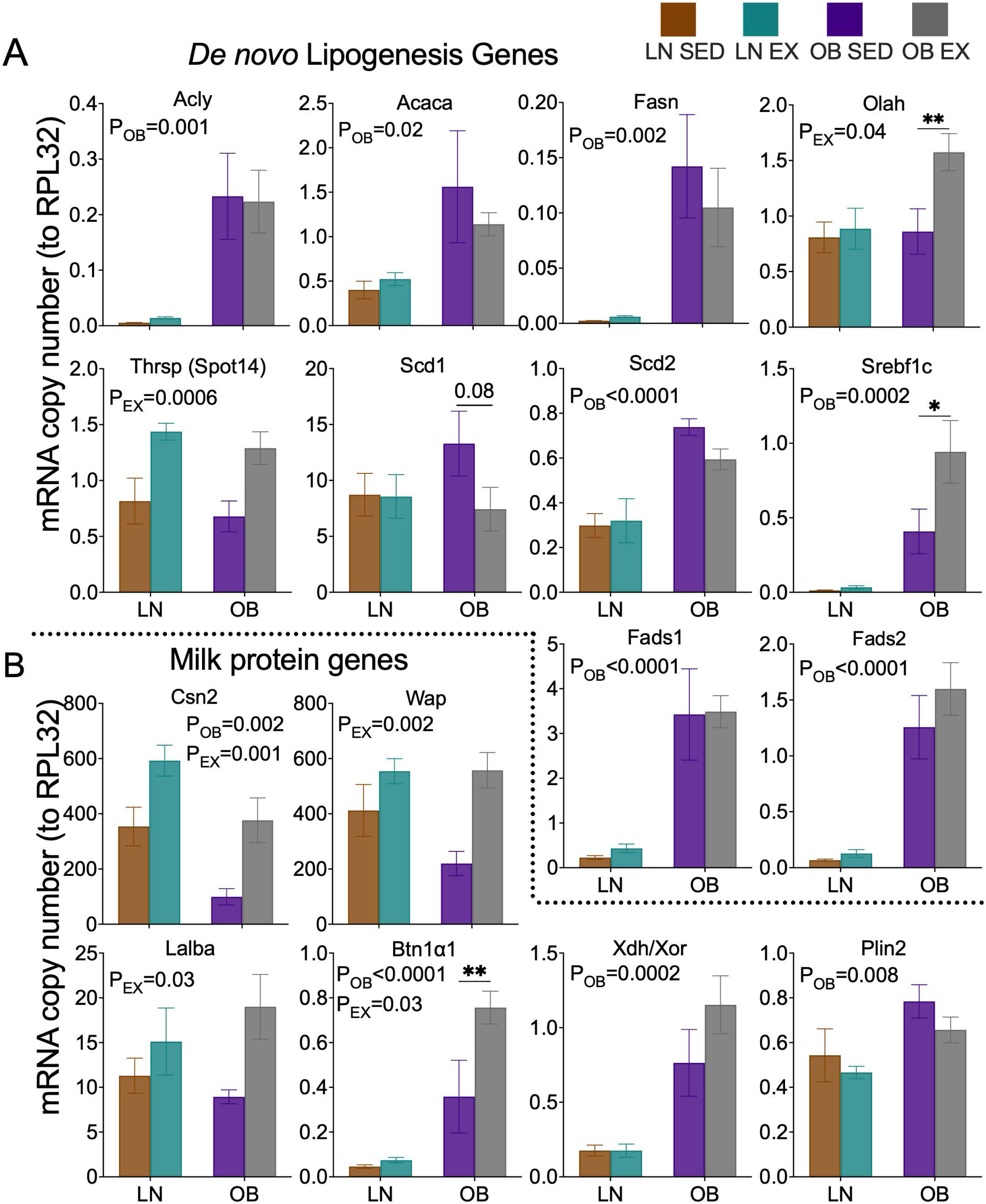

